# Motor selection dynamics in FEF explain the reaction time variance of saccades to single targets

**DOI:** 10.1101/221218

**Authors:** Christopher K. Hauser, Dantong Zhu, Terrence R. Stanford, Emilio Salinas

## Abstract

In studies of voluntary movement, a most elemental quantity is the reaction time (RT) between the onset of a visual stimulus and a saccade toward it. However, this RT demonstrates extremely high variability, which in spite of extensive research remains unexplained. It is well established that, when a visual target appears, oculomotor activity gradually builds up until a critical level is reached, at which point a saccade is triggered. Here, we further characterize the dynamics of this rise-to-threshold process based on computational work and single-neuron recordings from the frontal eye field (FEF) of behaving monkeys. We find that the baseline activity, build-up rate, and threshold level show strong, nonlinear co-dependencies that explain the distinct RT distributions observed experimentally. The results indicate that intrinsic randomness contributes little to saccade variance, which results mainly from an intricate, fundamentally deterministic mechanism of motor conflict resolution that has subtle yet highly characteristic manifestations.

---

The reaction time **(RT)** represents the total time taken by all the mental operations that may contribute to a particular action, such as stimulus detection, attention, working memory, and motor preparation. While its importance as a fundamental metric for inferring the mechanisms of both normal and pathological brain function cannot be overstated (Welford, 1980; Meyer et al., 1988), such reliance is a double-edged sword. Under appropriate experimental conditions, differential measurements of RT may be used as a readout for changes in the (mean) time consumed by any one of the aforementioned operations. However, a particular RT value is hard to interpret because not all the operations may be known, and those that are relevant may overlap in time to varying degrees. Furthermore, each operation may have its own, independent source of variability, making it very difficult to attribute the measured variance in RT to a particular cause (e.g., Krajbich et al., 2015). We believe the study of saccadic eye movements is singularly susceptible to this type of confound. There is a seemingly firm, mechanistic account of what determines the variability of saccadic RTs — but according to the present results, that account is both incomplete and, in many ways, incorrect.

The saccadic system is widely used not only as a model for studying motor control but also as the output stage of a wide variety of decision-making processes (often assessed via RT measurements) (Liversedge et al., 2011). Hence, the neural dynamics that give rise to eye movements are well established. In essence, the preparation to make a saccade of a particular direction and amplitude is equal to a gradual rise in the activity of oculomotor neurons that are selective for the corresponding movement vector. If this rising activity, referred to as a motor plan, ramps up rapidly, the saccade is initiated soon; if the motor plan grows more slowly, the saccade starts later. Quantitatively, this corresponds to a negative correlation between saccadic RT and build-up rate (Hanes and Schall, 1996; Fecteau and Munoz, 2007; Ding and Gold, 2012; Heitz and Schall, 2012; Costello et al., 2013). Notably, neurons that encode motor plans seem to reach a consistent firing level just before the onset of a saccade, particularly in the superior colliculus **(SC)** and the frontal eye field **(FEF)** (Hanes and Schall, 1996; Brown et al., 2008; Stanford et al., 2010; Ding and Gold, 2012). That is, there appears to be a fixed activity threshold that serves as a trigger for eye movements (Lo and Wang, 2006). Thus, it is widely thought that, for simple saccades to lone, unambiguous stimuli, or reactive saccades, the variance of the RT distribution is predominantly determined by the variance of the FEF/SC build-up rates across trials (Carpenter and Williams, 1995; Hanes and Schall, 1996; Fecteau and Munoz, 2007; Sumner, 2011).

This rise-to-threshold process is of enormous conceptual importance, as it is the key buildingblock of virtually all models of decision making in which multiple choice alternatives, typically guided by perceptual information, are evaluated over time (Gold and Shadlen, 2001; Erlhagen and Schoner, 2002; Smith and Ratcliff, 2004; Brown and Heathcote, 2008; Stanford et al., 2010; Krajbich and Rangel, 2011; Thura et al., 2012; Brunton et al., 2013; Miller and Katz, 2013). However, the variance of this process in its simplest possible instantiation — reactive saccades — remains a mystery (Sumner, 2011), because it seems too large to just reflect noise, or intrinsic randomness (in the build-up rates of oculomotor neurons). One possibility is that the randomness is purposeful, that unpredictability in saccade timing somehow entails a behavioral advantage (Carpenter, 1999). Alternatively, the RT of reactive saccades may fluctuate at least in part because of underlying neural mechanisms that have simply not been identified yet.

**Figure 1.**
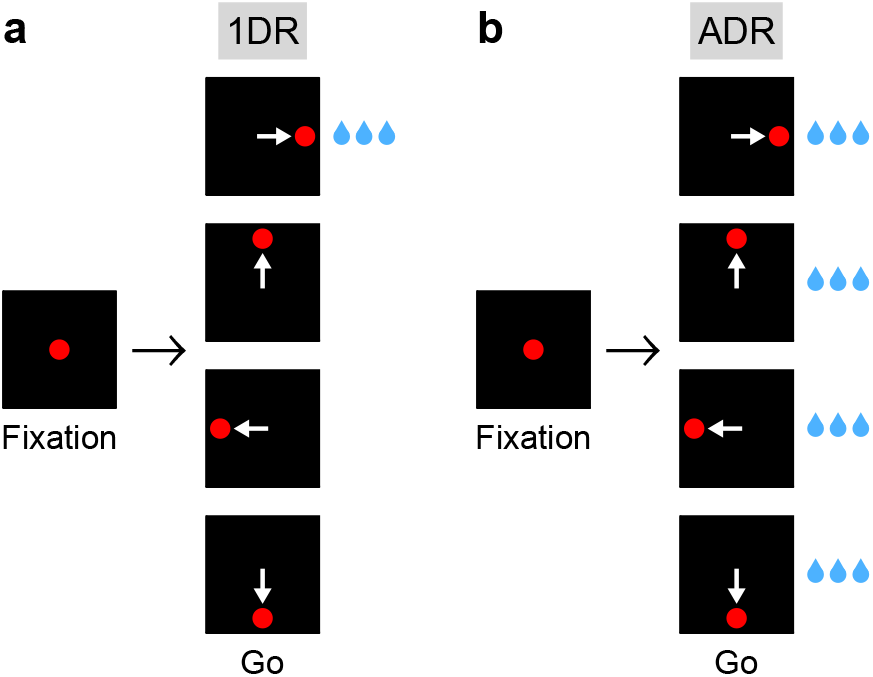
Saccadic tasks used. **a**, The 1DR task. After a fixation period of 1000 ms, a single eccentric stimulus appears at one of four locations and the subject is required to make a saccade to it, Stimulus location is chosen randomly in each trial. Fixation offset (go signal) and stimulus onset are simultaneous, In each block of trials, only one of the directions yields a large reward; the others yield either no reward (monkey G) or a small reward (monkey K). **b**, The ADR task. Same sequence of events as in **a**, except that saccades in all directions are equally rewarded. White arrows indicate saccades; they are not displayed.

Here, we describe such mechanisms. We recorded activity from single FEF neurons in an elegant paradigm (Lauwereyns et al., 2002; Hikosaka et al., 2006) that produces a large spread in saccadic RT simply by varying the subject's spatial expectation of reward. As anticipated, we saw that the build-up rate of the rise-to-threshold process is inversely related to RT, but we also found that the variations in build-up rate across trials are strongly linked to large fluctuations in both threshold and baseline activity (measured during fixation). A model based on competitive dynamics quantitatively reproduced the temporal profiles of the evoked neural responses, as well as their dependencies on RT, reward expectation, and trial outcome (correct/incorrect) — this, while simultaneously matching the monkey's full RT distributions across experimental conditions. The results indicate that the noise in the build-up rate is much more modest than predicted by extant frameworks, and that the high variability of saccades to single targets is, to a large degree, deterministic, a direct consequence of motor selection mechanisms that allow voluntary saccades to be driven by both sensory events and internal biases.

## Results

### Behavioral manifestations of a spatial bias

Two rhesus monkeys were trained on the one-direction rewarded **(1DR)** task (Fig. 1a), in which a saccade to a lone, unambiguous target must be made but a large liquid reward (primary reinforcer) is available only when the target appears in one specific location (Lauwereyns et al., 2002; Hikosaka et al., 2006). The rewarded location remains constant over a block of trials and then changes. Of note, ours is a RT version of the task whereby the go signal meaning “move now!” (offset of fixation point) is simultaneous with target onset. Also, it involves 4 locations and variable block length. This task generates errors and a large spread in RT (Fig. 2) under minimalistic sensory stimulation conditions. We exploit this to investigate how variance in saccadic performance relates to variance in FEF activity.

When the target and rewarded locations coincided (congruent trials; Fig 2a, red traces), the monkeys consistently moved their eyes very quickly (monkey G, 158 ± 33 ms, mean RT ± 1 SD; monkey K, 146 ± 21 ms), and essentially never missed (monkey G, 99.8% correct, *n* = 7234 congruent trials; monkey K, 99.6%, *n* = 5837). In contrast, when the rewarded and target locations were either diametrically opposed or adjacent (incongruent trials; Fig. 2b-d, cyan traces), both the mean RT and the spread increased dramatically (monkey G, 269 ± 84 ms; monkey K, 236 ± 77 ms), along with the percentage of incorrect saccades away from the target (monkey G, 18.3% incorrect, *n* = 16905 incongruent trials; monkey K, 8.1%, *n* = 12708). The symmetric condition in which all directions were equally rewarded (**ADR**; Fig. 1b) produced RT distributions that were intermediate between those of congruent and incongruent trials (Fig. 2a-d, gray; monkey G, 192 ± 40 ms; monkey K, 174 ± 36 ms). These results recapitulate the puzzle mentioned in the Introduction: if saccades can be very fast, why, under identical stimulation conditions, are they sometimes very slow?

**Figure 2.**
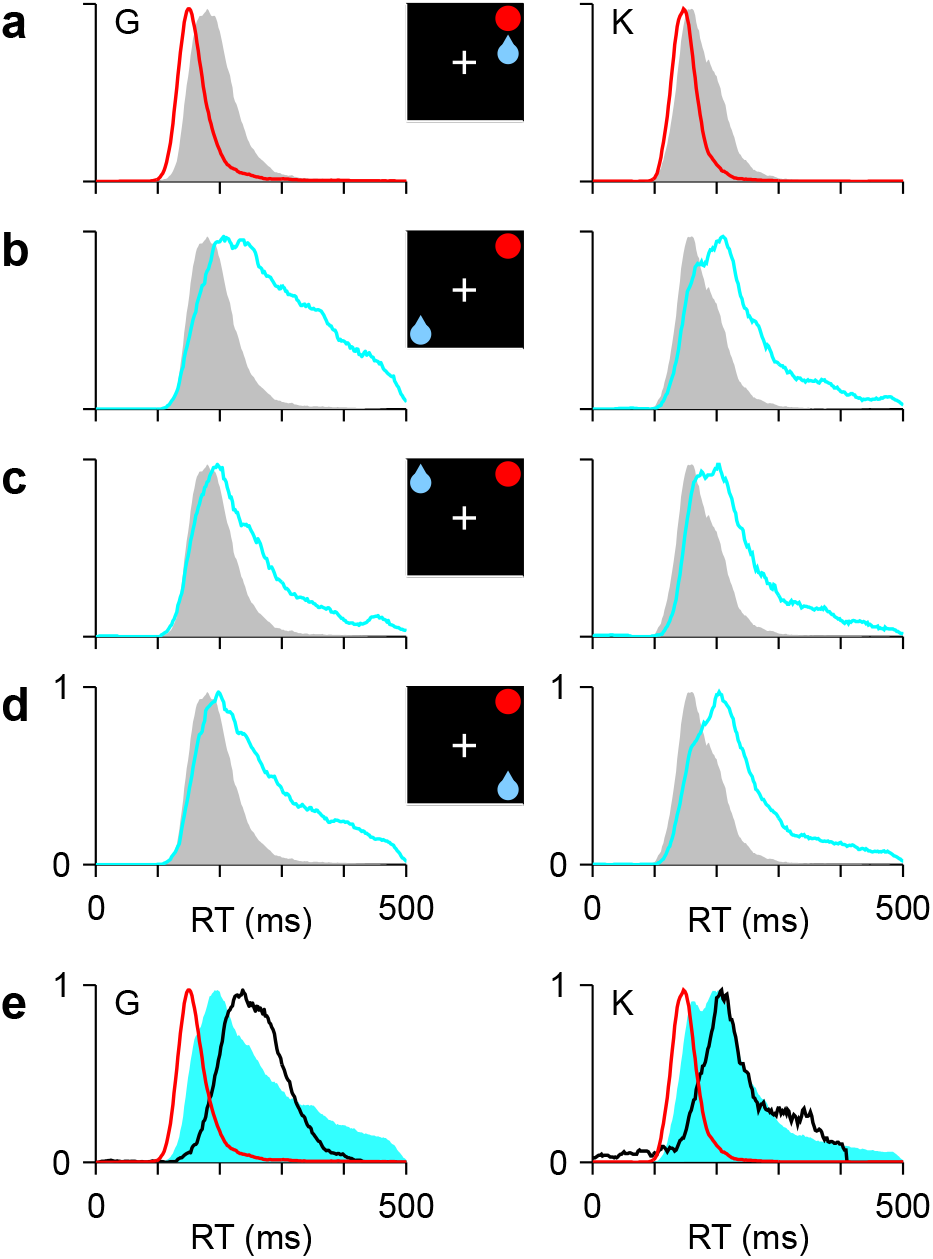
Asymmetric reward expectation leads to strong spatial bias. **a-d**, RT distributions for correct saccades, for monkeys G (left column) and K (right column). Insets indicate rewarded location (blue drop) and target (red circle). When the two are congruent (**a**, red traces), RTs are shorter and less variable than when they are incongruent, i.e., either opposite (**b**, cyan) or adjacent (**c, d**, cyan). In unbiased trials (ADR task; gray), results are intermediate. **e**, RT distributions in correct congruent (red, same data as in **a)**, correct incongruent (cyan, data in **b-d** combined), and incorrect incongruent (black) trials. Histograms are normalized to a maximum of 1. The RTs during errors are neither the fastest nor the slowest.

Also note that, compared to those of correct saccades, the RTs of incorrect saccades were neither consistently fast, as might be expected based on strong anticipation, nor consistently slow, as might be expected from a protracted conflict resolution process (Fig. 2e). Instead, the RTs during errors fell squarely in the middle of the distributions of correct RTs (for correct trials 90% of RTs were inside [158, 432] and [146, 404] ms for monkeys G and K, respectively; for errors, the ranges were [180, 336] and [152, 388] ms). Mechanistically it is not obvious how this could be accomplished. One of the aims of the model presented below is to reproduce the RT data shown in Fig. 2 and resolve this conundrum.

In short, monkeys are highly sensitive to the spatially asymmetric value associated with otherwise identical target stimuli (for additional evidence of this, see Supplementary Fig. 1). This manifests as large differences in RT across three experimental conditions: correct congruent, correct incongruent, and incorrect incongruent trials. The pattern of results suggests that, when a target appears at an unrewarded location, a conflict arises that leads to high variability in the direction and timing of the elicited saccade. In what follows, we ask: what features of the evoked FEF activity may reflect that conflict and account for the variability of the measured RTs?

### Motor conflict during the rise-to-threshold process

We recorded single-unit activity from 132 FEF neurons in the two monkeys (67 in monkey G; 65 in monkey K) during performance of the 1DR task. The analysis in this section focuses on a population of 62 neurons that satisfied two conditions: they had standard visual and saccade-related responses (for details of the cell classification procedure, see Methods), and both correct and incorrect trials were collected for them.

In this section we show that, during the rise-to-threshold process associated with reactive saccades, the build-up rate, the apparent threshold, and the baseline firing rates measured during fixation (before the target is presented) exhibit coordinated variations that have not been appreciated before and which, beyond the specifics of our task, are likely to be key for determining RTs in general. We refer to the baseline levels in plural because the target location is not the only relevant one; any internally-driven bias favoring other locations to which the saccade may be directed creates motor conflict, i.e., activity that competes with and impacts the target-driven response — and the RT.

Before describing these results, a note about nomenclature. Whereas just 2 distinct experimental conditions, congruent and incongruent, are relevant for behavioral analysis, neurophysiologically there are 8 conditions to consider, depending on whether the target, expected reward, and saccadic movement were inside or outside a recorded cell's response field (**RF).** Thus, for example, we use the abbreviation IOI to denote the target/reward/saccade combination in/out/in; that is, trials for which the target was in, the reward was expected out, and the saccade was into the RF (see icons in Fig. 3a-c). Such IOI trials are incongruent, because target and reward locations do not match, and correct, because target and saccade locations match. Only 6 of the 8 possible combinations are considered because congruent trials were virtually devoid of errors, so IIO and OOI conditions are absent. With this notation at hand, we now turn to the activity of FEF across those 6 remaining conditions.

Although each saccade in the 1DR task involves a single target, the motor preparation process in FEF can (and should) be understood as a motor selection process involving not one, but at least two populations of neurons, those that contribute to the actual saccadic choice and those that favor the opposite choice (Fig. 3a-c, see icons). During congruent trials, when the target appears at the rewarded location, the evoked activity rises most rapidly and reaches the highest firing rate (Fig. 3a, III trials, red trace). The saccade is essentially always correct, and no evidence of conflict is discernible, as the neurons favoring the opposite choice, away from the target, show little, if any, response before the eye movement (Fig. 3a, OOO trials, green trace). In this case, naively, it would appear as if only the one response that rises to threshold is important.

Notably, though, a difference in activity between the two complementary populations is already evident before the go signal/target onset. That is the internal bias signal created by reward expectation, which in this case is spatially congruent with both the target and the saccade. This baseline signal is consistent with previous neurophysiological studies using the 1DR task (Takikawa et al., 2002; Sato and Hikosaka, 2002; Ikeda and Hikosaka, 2003), and may be interpreted as a neural correlate of spatial attention (Maunsell, 2004; Preciado et al., 2017).

By contrast, during incongruent trials, when the target is presented opposite to the rewarded location, a conflict arises early in the trial in the form of a higher baseline favoring the rewarded location (Fig. 3b, note green trace above red before go signal). During correct trials this conflict is appropriately resolved as the target-driven activity increases and overtakes the competition (Fig. 3b, IOI trials, red trace), but the rise proceeds more slowly, i.e., it has a lower build-up rate, and ultimately reaches a lower peak firing level (*R_p_*; see Methods) than that observed when the bias and the target are congruent (Fig. 3d, IOI vs. III). Finally, the conflict is even more extreme during incongruent trials that end in erroneous choices toward the rewarded location (Fig. 3c). In that case the initial bias in baseline activity is largest (Fig. 3e, IOO vs. OII), and the evoked target-driven activity (Fig. 3c, magenta trace) is considerably weaker compared to that observed during correct saccades (Fig. 3d, IOO vs. IOI). The neural response associated with the (wrong) saccadic choice rises very slowly (Fig. 3c, cyan trace) and reaches a modest threshold level prior to saccade onset, but this activity is nonetheless slightly above that associated with the opposite (correct) motor alternative (Fig. 3c, right panel; Fig. 3d, OII vs. IOO). This ambivalence between motor plans is reminiscent of that observed during choice tasks (Thompson et al., 2005; Ding and Gold, 2012; Costello et al., 2013), as if the monkeys had struggled to make a choice between two competing targets, even though only one was displayed.

**Figure 3.**
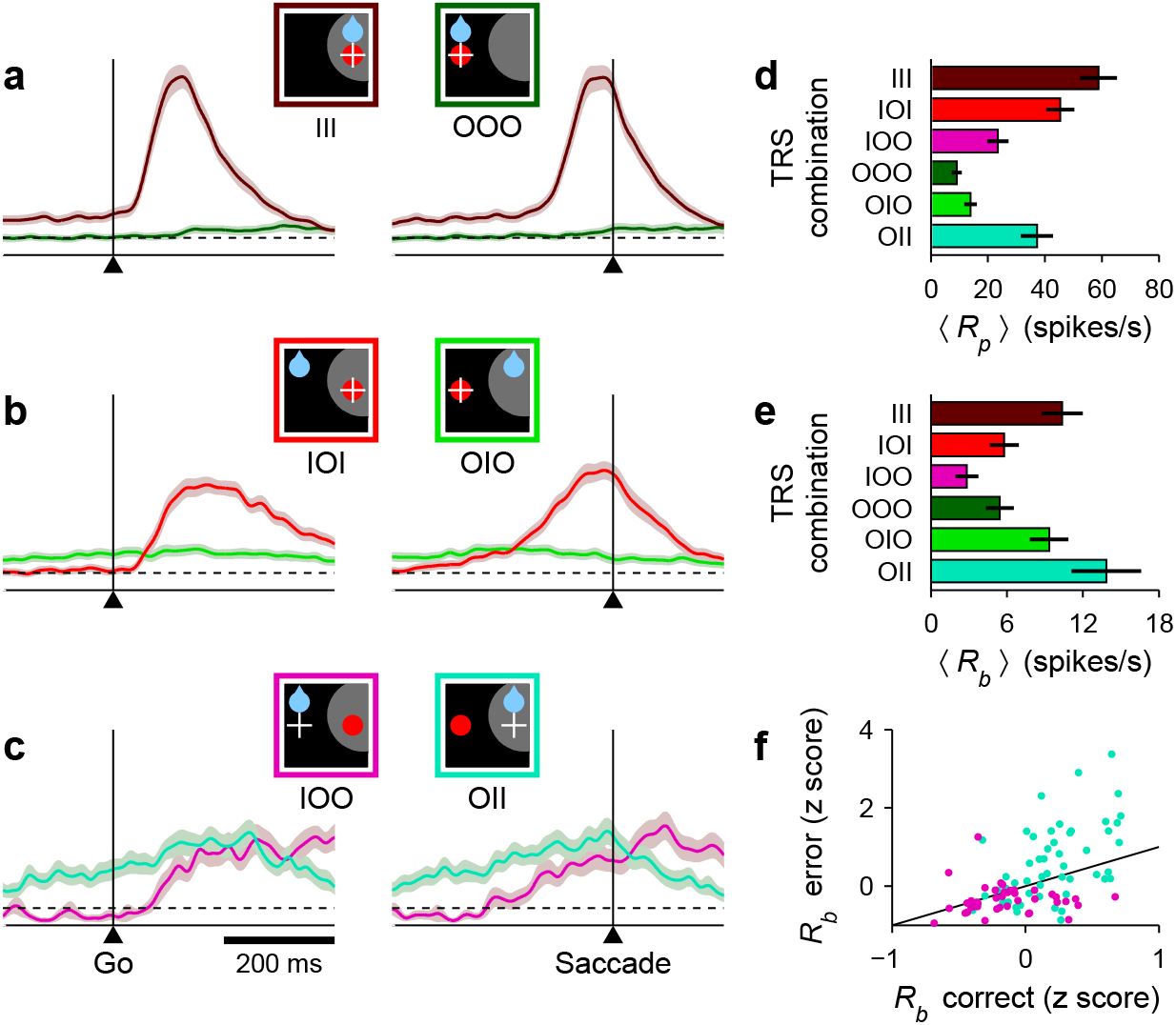
Baseline activity predicts response gain, threshold, and outcome. **a-c**, Normalized firing rate as a function of time for a population of 62 neurons (V, VM, and M) for which both correct and incorrect responses were collected. Icons indicate target (red circle), rewarded location (blue drop), and saccade (white cross) relative to the RF (gray area) in each case. Paired reddish and greenish traces correspond to activity with the target inside or outside the RF, respectively, in the same behavioral condition. For congruent trials (**a**) only correct responses are shown. For incongruent trials both correct (**b**) and incorrect (**c**) responses are shown. The reference line (dotted) is identical across panels. **d, e**, Mean peak activity (**d**) and mean baseline activity (**e**) for each target/reward/saccade (TRS) combination, from the same 62 neurons in **a-c.** Error bars indicate ±1 SE across cells. **f**, Baseline activity (z-scored) for incorrect (y axis) versus correct outcomes (x axis) under identical target/reward conditions. Each point is one neuron. Magenta dots indicate IOO versus IOI trials (target in/reward out); cyan dots indicate OII versus OIO trials (target out/reward in).

These results were based on recordings from 62 neurons with diverse visuomotor properties, but were qualitatively similar when the averaging across cells was restricted to units that were either predominantly visual (Supplementary Fig. 2a-c) or predominantly motor (Supplementary Fig. 2d-f). Those two populations differed in the time at which they fired maximally, but their responses were otherwise similar across conditions and outcomes. In both cases the motor preparation process during errors was characteristically different from that during congruent trials: the activity build-up was more gradual and reached a lower threshold, and the opposing motor plans were much more ambivalent.

That expectation (and/or attention; Maunsell, 2004; Preciado et al., 2017) is what drives the variations in baseline is clear; during each block, the rewarded location is known to the subject during the fixation period, whereas the target location is not. So it is important to underscore that, during incongruent trials, the baseline activity *R_b_* (firing rate in a 250 ms window preceding target onset) is strongly predictive of outcome. In correct trials, *R_b_* was higher for the rewarded location than for the opposite (target) location (Fig. 3e, OIO vs. IOI; *p* < 10^−5^, permutation test), but such difference was particularly large during incorrect saccades (Fig. 3e, OII vs. IOO; *p* = 10^−4^). These variations in activity were consistent across a majority of individual cells. When the rewarded location coincided with the RF and the target was presented outside, most neurons (34 of 53, *p* = 0.03, binomial test) had a higher baseline rate before incorrect as opposed to correct trials (Fig. 3f, cyan dots, OII vs. OIO; *p* = 0.002, permutation test), as if an excessive *R_b_* triggered a (wrong) saccade into the RF. Conversely, when the rewarded location was opposite to the RF and the target was subsequently presented inside, most neurons (33 of 42, *p* = 0.001, binomial test) had a lower *R_b_* preceding incorrect trials (Fig. 3f, magenta dots, IOO vs. IOI; *p* = 0.009), as if the lack of baseline activity precluded a (correct) saccade into the RF.

In summary, during the 1DR task we observed characteristic variations in the three key features of the rising saccade-related activity in FEF: the ‘baseline’ firing rates at the target and opposite spatial locations, the build-up rate of the target-driven response, and the apparent threshold reached before movement onset. All three quantities, together with the degree of conflict between the opposing motor plans, varied depending on congruency and outcome. Notably, the baseline signal arises earliest, before target presentation, and is highly predictive. Next, we show that these novel interrelationships provide key constraints for a new mechanistic model that relates neuronal activity to RT with remarkable detail.

### Modeling the rise-to-threshold dynamics

A saccadic competition model was developed to investigate the mechanistic link between the FEF activity and the monkeys’ behavior in the 1DR task (Methods). Such a bridge requires that multiple constraints be satisfied. First of all, the model must reproduce the neurophysiological results presented in the previous section. Thus, it considers two neural populations whose responses may rise to a threshold. One population generates saccades toward location *T*, where the target stimulus is presented, and the other toward D, the diametrically opposite location (Fig. 4a, icon). In any given trial, if the target-driven response, *R_T_* (Fig. 4a-c, red traces), reaches threshold first, a correct saccade to the target is produced, whereas if the bias-driven activity, *R_D_* (Fig. 4a-c, blue traces), reaches threshold first, the result is an incorrect saccade away from the target. In this way, the *R_T_* and *R_D_* variables correspond to the population responses recorded with the target inside (Fig. 3a-c, reddish traces) and outside (greenish traces) of the RF, respectively.

Another important feature of the recorded data is the evident asymmetry between the two motor plans: the target-driven activity is typically strong and never fully suppressed, whereas the internally-driven activity favoring the opposite location is typically suppressed and only rarely of moderate strength. In the model, this asymmetry is captured by two suppression mechanisms that constrain when and how *R_D_* can rise. One of them (Methods, Rule 1) simply prescribes that, once *R_T_* is rising, it can fully suppress *R_D_*. That is, the moment *R_T_* advances past *R_D_*, *R_D_* stops rising altogether. The other mechanism is about the timing of *R_D_*. Whereas the target-driven response, *R_T_*, starts rising shortly after target onset (after an afferent delay of 35 ms), its counterpart, *R_D_*, starts rising later, partly because of a somewhat longer afferent delay (50 ms) but mostly because of a transient, stimulus-driven suppression. This (partial) suppression is based on abundant evidence indicating that ongoing saccadic plans, *R_D_* in this case, are briefly inhibited by stimuli that appear abruptly, just like the target in our experiment (Reingold and Stampe, 2002; Dorris et al., 2007; Stanford et al., 2010; Bompas and Sumner, 2011; Buonocore and McIntosh, 2012; Buonocore et al., 2017). In the model, the inhibition lasts 115 ms (Fig. 4a-c, gray shades), after which *R_D_* may rise in full force — if it was not overtaken by *R_T_* in the interim. These two suppression mechanisms give the target-driven activity an advantage over its competing, internally-driven counterpart, and are necessary to account for the late-onset, weak activity away from the RF seen during correct saccades (Figs. 3a, b, greenish traces). Notably, the suppression mechanisms are also consistent with the behavior; without them, the model would produce incorrect saccades at a vastly higher rate than observed experimentally and with an exceedingly high proportion of fast RTs.

**Figure 4.**
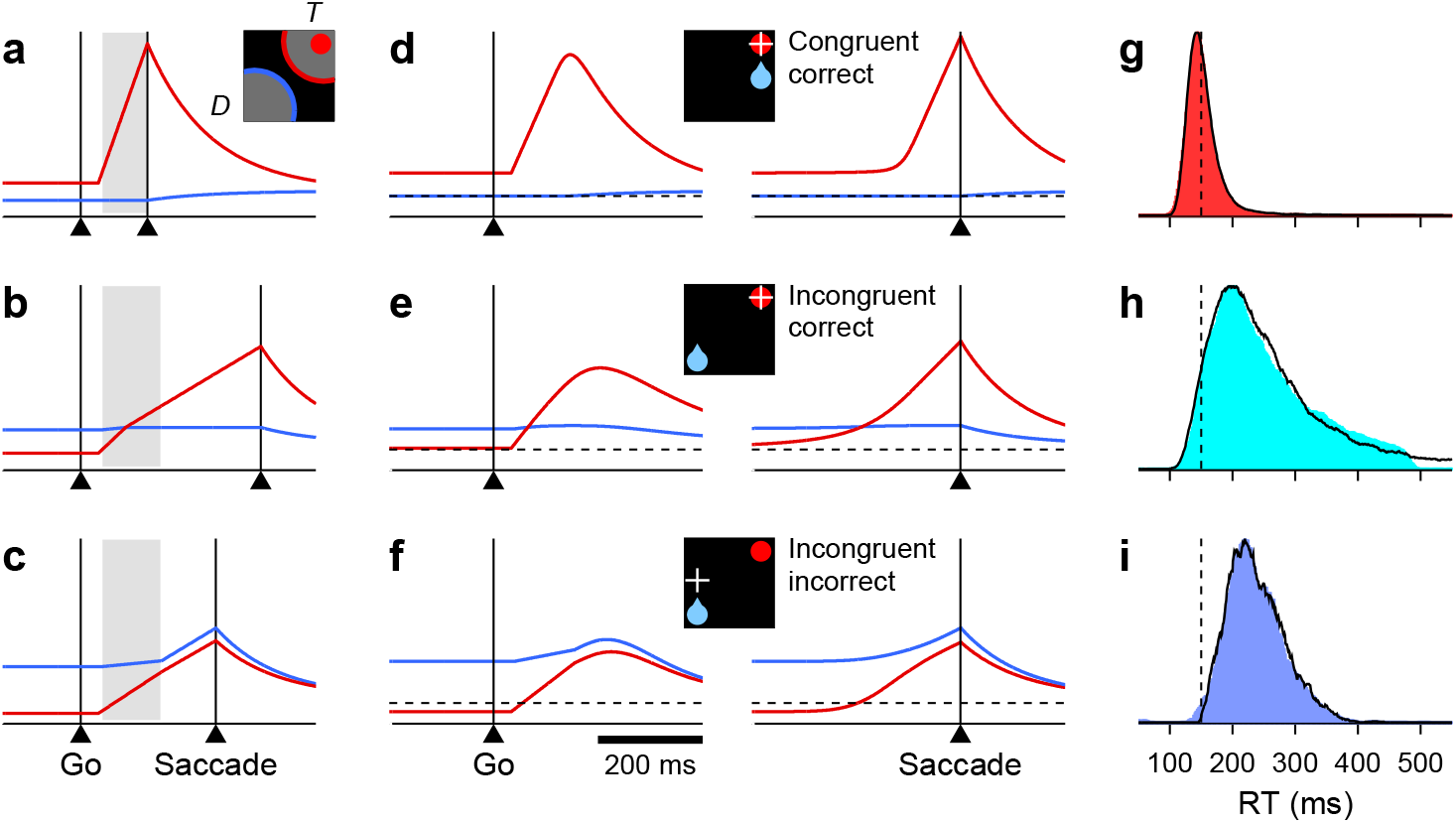
Simulation results from a saccadic competition model that bridges the neural and behavioral data. **a-c**, Simulated activity in three single trials. Traces show motor plans toward the target location (*T*, red) and the diametrically opposite location (D, blue). Gray shades mark the period during which the target stimulus suppresses the motor plan toward D. When the target-driven activity, *R_T_* (red), reaches threshold first (**a, b**) the saccade is correct; when the bias-driven activity, *R_D_* (blue), reaches threshold first (**c**) the saccade is incorrect. The scale on the y axes is the same for all three plots. **d-f**, Mean firing rate traces *R_T_* (red) and *R_D_* (blue) averaged across correct congruent (**d**), correct incongruent (**e**), and incorrect incongruent (**f**) simulated trials. Same format as in Fig. 3a-c. **g-i**, RT distributions from the same simulations as in panels **d-f**, with trials sorted accordingly: for correct congruent (**g**), correct incongruent (**h**), and incorrect incongruent (**i**) trials. Colored shades are combined data from the two monkeys; black lines are model results. Dotted vertical lines at 150 ms are for reference.

Finally, the model must also capture the variations in baseline activity, build-up rate, and threshold observed in the FEF data (Fig. 3d, e), and this is where the crucial conceptual leap takes place. What we found empirically was that, for the target-driven response, a higher baseline was accompanied by both a higher build-up rate and a higher threshold (with the baseline activity of the alternative motor plan having opposite effects). The model generalizes these dependencies by making two assumptions. First, that similar relationships hold across *all trials*, rather than just across the three experimental conditions examined, and second, that because the baseline signal is present before target onset, any variations in build-up rate and threshold can be formulated mainly as the result of variations in baseline activity. Thus, the model can be thought of as designed to test whether the differences in the rise-to-threshold process observed across experimental conditions (Fig. 3) are the average manifestations of similar but more general dynamical relationships between the three relevant variables, where the variance is primarily derived from the baselines. So, in practice, the general idea is that the baselines fluctuate stochastically and determine the ensuing rise-to-threshold excursion in each trial.

The resulting dynamics between competing motor plans can be intuitively appreciated with three example trials (Fig. 4a-c). The simplest situation is when, during fixation, the baseline at the target location, *B_T_*, is larger than that at the opposite location, *B_D_* (Fig. 4a). This is typically the case when the target and rewarded locations coincide (but note that what matters in the model is simply the actual baseline values; more on this below). The condition *B_T_* > *B_D_* has two specific consequences: (1) it produces a high build-up rate for the target-driven activity (Equation 7), *R_T_* (red trace), and (2) it sets the saccade threshold, Θ, to a high value (Equation 6). Thus, because of the high build-up rate, *R_T_* rises sharply and quickly triggers a saccade, in spite of the high Θ. The *D* plan (blue trace) is always suppressed in this case, so no overt conflict is visible. This is how correct saccades with very short RTs are produced.

The more interesting scenario occurs when the bias-driven plan starts with the higher baseline, as typically happens when the reward is expected at the *D* location (but again, the dynamics are dictated just by the baseline values). In that case the saccade can be either correct or incorrect, depending on how big the lead is. When *B_D_* is much larger than *B_T_* (Fig. 4c), the consequences are essentially the opposite of those in the previous example: (1) *R_T_* has a low build-up rate, so the target-driven response (red trace) rises slowly, and (2) the saccade threshold, Θ, is low. In this way, *R_D_* is able to advance steadily after the suppression interval and win the race from wire to wire, reaching a relatively low firing level before saccade onset. This is how incorrect saccades are produced.

In contrast, if the baseline *B_D_* is only moderately higher than *B_T_* (Fig. 4b), then the combined effect of the baselines is intermediate relative to that of the two previous examples: (1) the (initial) build-up rate of *R_T_* is moderate, neither as high as in the first example nor as low as in the second, and (2) the value of Θ is also intermediate. Thus, the target-driven plan (red trace) rises at a rate that allows it to overtake the competing plan (blue trace) and win the race coming from behind. Importantly, in this case *R_T_* slows down as it goes past *R_D_* (note slight change in slope of red trace during shaded interval). Although *R_T_* wins the race, overtaking the competing plan exacts a toll, and the lower its initial build-up rate, the higher that toll (Equation 11). This is the one mechanism that was introduced in the model specifically to satisfy key behavioral constraints. In this case, slowing down the *winner* target-driven plan is necessary for producing correct saccades with very long RTs — longer than those of incorrect saccades.

These examples illustrate how the baseline levels *B_T_* and *B_D_* quantitatively regulate both the build-up rate of the target-driven activity and the saccade threshold, but it is important to stress that, at the same time, the baselines convey the information about the location of the expected reward in a manner that is consistent with the experimental data. In the simulations, the fluctuations of the baselines across trials are characterized by their mean and variance. The variance is determined by a single, fixed parameter (*σ*; see Methods). The two mean baseline values are also fixed, but are assigned to the *T* and *D* locations according to a simple prescription: the rewarded location gets the higher mean (Equation 5). Thus, in simulations of the congruent case *B_T_* is, on average, larger than *B_D_* (as in Fig. 4a), and the majority of trials are fast and correct; whereas in simulations of the incongruent case the roles are reversed, *B_T_* is, on average, lower than *B_D_* (as in Fig. 4b, c), which results in a combination of correct and incorrect slower responses. Other than that, the simulations proceed in exactly the same way in the two bias conditions, as they should.

In this way, when the simulated firing rate trajectories are sorted by bias and outcome, the model reproduces the covariations in baseline, build-up rate, and threshold across conditions (Fig. 4d-f; for quantification, see Supplementary Fig. 3). This demonstrates that, as intended, the hypothesized coupling across trials is indeed consistent with the observed coupling across experimental conditions. In addition, the average *R_T_*(t) and *R_D_* (t) traces match the trajectories of the recorded population responses, as well as the varying degree of ambivalence between the two motor alternatives, in great detail (compare to Fig. 3a-c). The proposed interaction mechanisms between the two motor plans result in average traces with the appropriate magnitude and time course. But most critically, at the same time, the model fully accounts for the behavioral data: (1) it generates correct and incorrect saccades in proportions similar to those found experimentally (∼0% and ∼10% errors in congruent and incongruent conditions), and (2) it generates simulated distributions of RTs (Fig. 4g-i) that closely mimic their behavioral counterparts (as assessed by mean, median, SD, and skewness). In particular, the RTs in incorrect trials (panel i) are neither too fast, because the stimulus-driven suppression mechanism prevents fast errors, nor too slow, because the slowest responses, which occur when *R_T_* slows down, are correct. The model explains the behavioral data in terms of the neural data, accurately replicating both.

According to the results, the baseline activity, build-up rate, and threshold vary in a coordinated fashion to generate the wide range of RTs observed in the task. In the rest of the paper we show that this fact explains many other, seemingly odd features of the neural data.

### Coupled variations in threshold and baseline

A key assumption of the model is that fluctuations in baseline activity result in fluctuations in build-up rate and threshold. To further dissociate the interdependencies between these three variables and, in particular, to determine whether baseline and threshold are directly coupled, we compared the responses evoked in congruent versus incongruent trials before and after equalizing their RTs.

The FEF responses recorded during III and IOI trials were quite distinct (Figs. 3a, 3b), even though both involved correct saccades in the same direction. The differences could be due to the different expected reward locations in the two conditions, which correlate most strongly with baseline activity, and/or to the different RTs generated (Fig. 5a), which depend most strongly on build-up rate. To eliminate the differences due to RT — and, presumably, to build-up rate — we devised a simple sub-sampling procedure (Methods). The idea is straightforward: the fastest IOI and slowest III trials are paired so that the resulting data subsets have identical numbers of trials and nearly identical RT distributions (Fig. 5b). Then we made comparisons across conditions with and without matching the RTs.

What should be expected based on the model? In the standard case, without RT matching (NM condition), the target-driven response, *R_T_*, has a higher baseline and reaches a higher threshold in congruent as compared to incongruent trials (Fig. 5c). However, because they have very different build-up rates (note steeper rise of magenta curve), aligned on saccade onset the corresponding response trajectories intersect each other twice. In contrast, when the mean and variance associated with RT are the same (YM condition), the shapes of the trajectories are much more similar (Fig. 5d). The residual effect, which is exclusively related to the internal bias signal, demonstrates somewhat more modest — but still clearly visible — differences in baseline and threshold, such that the response trajectory for congruent trials always stays above that for the incongruent (note magenta traces above black). These results make perfect sense within our modeling framework: first, because the build-up rate of the target-driven activity is strongly correlated with RT, so equalizing the RTs also equalizes the slopes of the neuronal trajectories; and second, because the coupling between the threshold and baseline across trials is also strong (Equation 6), so it remains clearly visible even when the RT range is drastically restricted.

**Figure 5.**
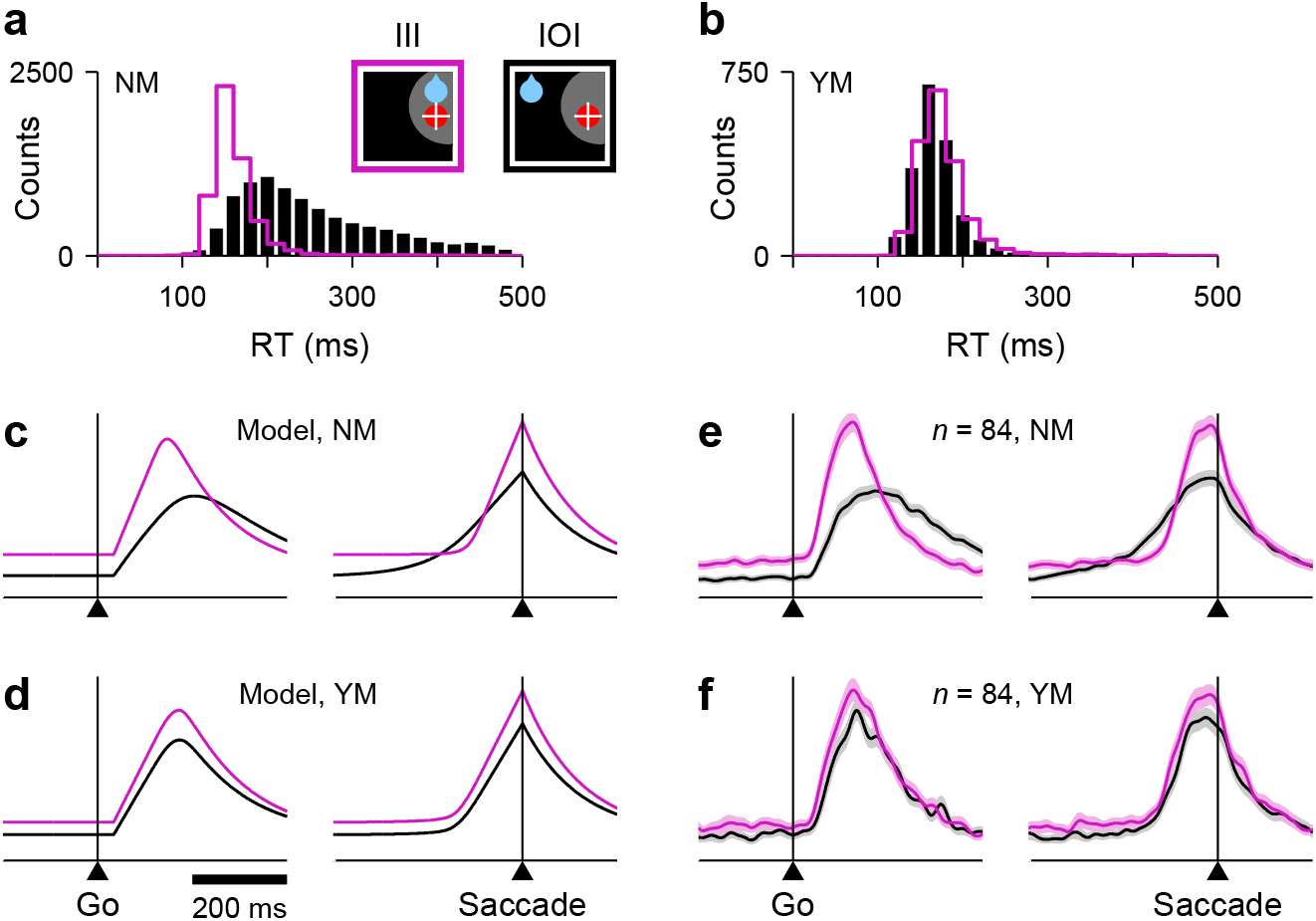
Disentangling the coupling between baseline activity, build-up rate, and threshold. **a**, Original RT distributions for III (magenta) and IOI (black) trials from all FEF recording sessions (non-matched condition, NM). **b**, RT distributions after RT matching (yes-matched condition, YM). **c**, Firing rate as a function of time for the target-driven activity, *R_T_*, in simulated III (magenta) and IOI (black) trials; same as red traces in Fig. 4d, e. RTs are not matched. **d**,As in **c**, but with matched RTs. **e**, Normalized firing rate as a function of time for a population of 84 V, VM, and M neurons. RTs are not matched. **f**, As in **e**, but with matched RTs.

Now consider the same analysis but for 84 V, VM, and M FEF neurons that had sufficient RT-matched trials. Qualitatively, the results are just as expected from the model: when the RTs are not matched, the responses in III and IOI trials differ patently in baseline, threshold, and build-up rate (Fig. 5e), and with the spikes aligned on saccade onset the corresponding curves intersect each other twice (right panel); in contrast, when the RTs are matched, the differences in build-up rate practically disappear, and the residual bias-related modulation is approximately equivalent to a vertical shift of the congruent (III) curve relative to the incongruent (IOI) one (Fig. 5f).

These results reveal the isolated effect of the reward-driven spatial bias. In addition, they lend further support to an important assumption of the model, namely, that the threshold for triggering a saccade is positively correlated with the baseline activity at the target location (as proposed earlier based on Fig. 3a, b). Such correlation exists regardless of the coupling of these variables to the build-up rate.

### Predicted RT sensitivity of the mean population activity

To test the model more stringently, we exploited the wide range of RTs produced in the 1DR task to generate predictions for how the evoked neural activity should be expected to vary as a function of RT. The rationale for these predictions is straightforward: instead of calculating the mean activity averaged across all trials in, say, the IOI condition (Fig. 4e), we consider, instead, similar traces based on subsets of trials within narrow RT bins (Methods). Assuming that the activity in FEF directly contributes to triggering each saccade, as happens in the model, the resulting response profiles should vary systematically across those RT bins, and any patterns should be consistent with the correlations between baseline, build-up rate, and threshold instantiated by the model, as well as with its other mechanisms (e.g., *R_D_* suppression).

Indeed, when the simulated IOI trials are sorted and averaged according to RT and the resulting curves are color-coded, a characteristic pattern emerges (Fig. 6c) in which steeply rising trajectories precede short-latency saccades (black) and more shallow, protracted trajectories precede long-latency saccades (red). This is largely because, in the model, the RT depends critically on the build-up rate of the target-driven activity. Notably, the apparent threshold reached by these curves at saccade onset is also modulated by RT (Fig. 6c, right panel), with fast choices (black) leading to higher activity levels than slow ones (red). This, in turn, is consistent with the correlated fluctuations in threshold and build-up rate instantiated in the model. This prediction for the IOI condition — i.e., the pattern resulting from the simultaneous dependencies of build-up rate and threshold on RT — reflects essential features of the model and, qualitatively, is highly robust to parameter variations.

**Figure 6.**
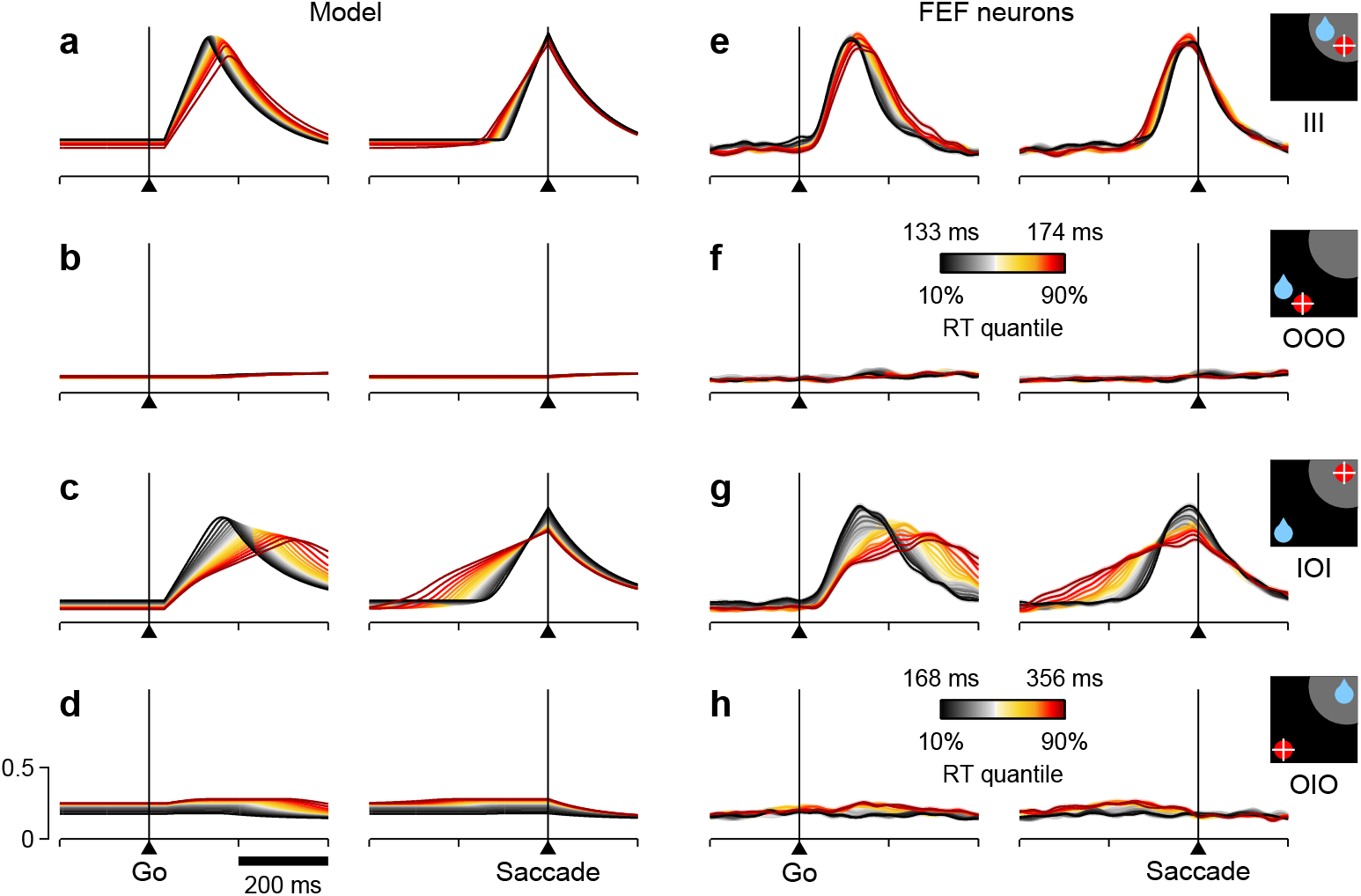
RT sensitivity of the average population activity. **a-d**, Firing rate as a function of time for the simulated motor plans *R_T_* (**a, c**) and *R_D_* (**b, d**) during congruent (**a, b**) and incongruent (**c, d**) trials. Equivalent experimental conditions are as indicated by the icons (far right). Curves are based on the same simulation runs as in Fig. 4d-f. Each colored trace includes 20% of the simulated trials around a particular *R_T_* quantile. **e-h**, Normalized firing rate as a function of time for a population of 84 FEF neurons (V, VM, and M). Activity is for correct saccades in the four experimental conditions indicated by the icons. Each colored trace includes 20% of the trials recorded from each participating cell around a particular RT quantile. Lighter shades behind lines indicate ± 1 SE across cells. Colorbars apply to both simulated and recorded data, either in congruent (III and OOO; colorbar in **f**) or incongruent conditions (IOI and OIO; colorbar in **h**). Scale bars in d apply to all panels.

**Figure 7.**
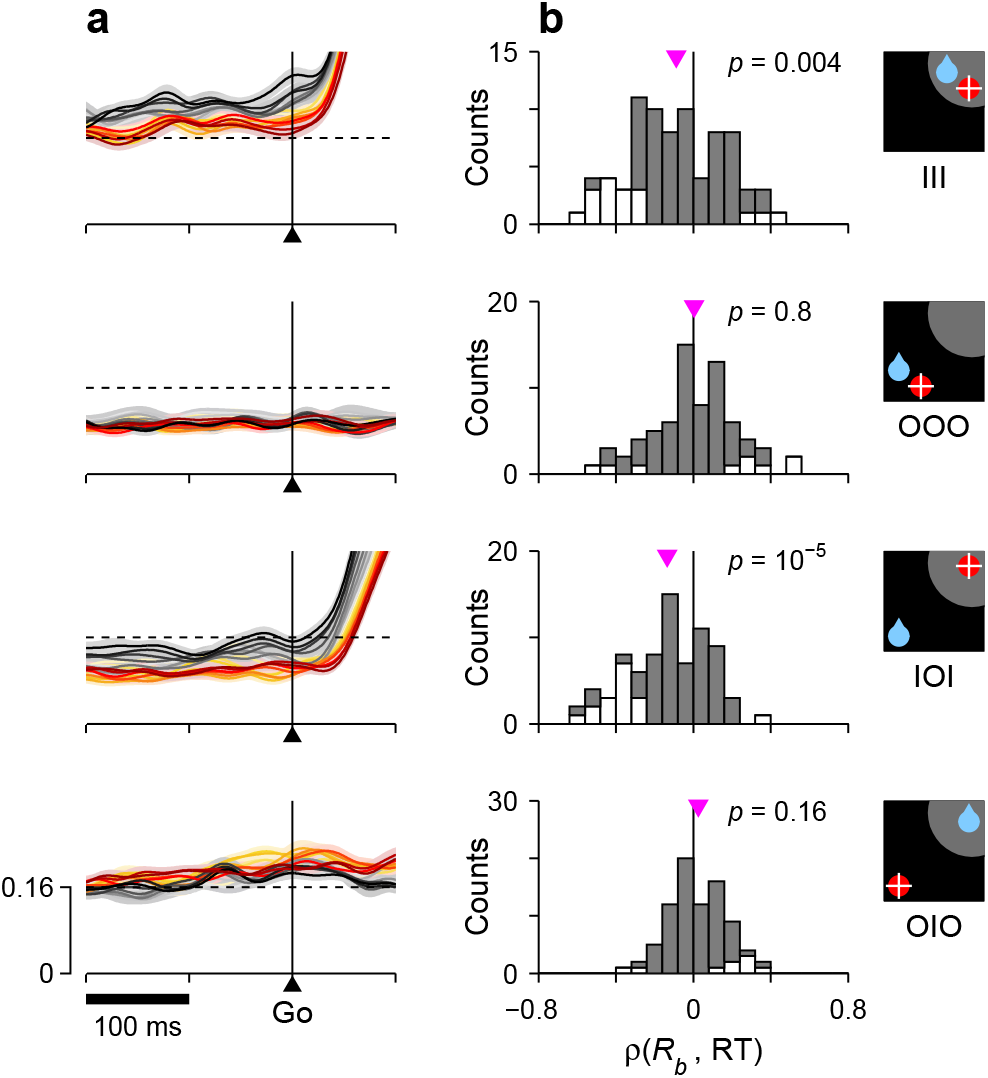
Correlation between RT and baseline activity in FEF. For each row, the corresponding target/reward/saccade configuration is depicted by the icon on the far right. **a**, Normalized neural responses around the go signal. Each colored trace corresponds to population firing rate as a function of time for a subset of trials around a particular RT quantile. Traces are exactly as in Fig. 6e-h (left panels), except at shorter x and y scales, as marked on the bottom panel. The dashed reference line is identical across panels. **b**, Distributions of Spearman correlation coefficients between RT and baseline activity, *ρ(R_b_*, RT). The data in each histogram are from the same 84 cells used in Fig. 6. White bars correspond to significant correlations (*p* < 0.05). Pink triangles mark mean values, with significance from signed-rank tests shown next to them.

To test this prediction, the recorded trials from 84 FEF neurons were sorted by RT in the same way as the simulated trials, and the corresponding traces were averaged across cells (Methods). The population curves that resulted (Fig. 6g) showed the same smooth transitions across RT bins as the simulated curves. Both the build-up rate and the threshold increased with shorter RTs as expected from the model.

More generally, when comparing across narrow RT bins, the agreement between the simulations and the overall population activity in FEF was always tight and evident — even though the model predictions varied widely across trial types. This was true in three important respects.

(1) For the activity evoked in III trials. When the target and rewarded locations were congruent, the neural responses into the RF (Fig. 6e) were much less sensitive to RT than for the corresponding incongruent trials (Fig. 6g), with the variations in threshold essentially disappearing (Fig. 6e, right panel). According to the model (Fig. 6a), this much weaker dependence resulted from intrinsic randomness, or noise, in the build-up rate, i.e, variability that is independent of the baseline (the term *η* en Equations 8, 9). In the model, such randomness is proportionally stronger in III than IOI trials, and blurs the effects created by the baseline-dependent fluctuations. This is explained in more detail in the next section. At the moment we simply emphasize that, although the sensitivity to RT manifested quite differently in III and IOI trials, there was close agreement between the simulated and neural data in both.

(2) For the baseline activity. The model postulates that the fluctuations in baseline translate into fluctuations in build-up rate and threshold. As a consequence, when sorted by RT, the simulated baseline levels spread accordingly. For the target-driven response, *R_T_*, a higher baseline always corresponds to shorter RTs (Fig. 6a, c, left panels, note black lines above red before go signal), so the correlation *ρ(R_b_*, RT) is negative. In contrast, for the baseline of the opposite motor plan, *R_D_*, the correlation is either positive (Fig. 6d, note red lines above black) or zero (Fig. 6b, note overlapping red and black curves), because of the competitive nature of the interactions between the *T* and *D* motor plans. Again, however, for all conditions the actual baseline activity measured in FEF was highly consistent with the model predictions — even though all the effects were expected to be small (Fig. 6e-h, left panels). This is easier to visualize when the data are magnified appropriately (Fig. 7a). In quantitative terms (Fig. 7b), on average, there was a significant negative correlation between baseline activity and RT in both III (*p* = 0.004, signed-rank test) and IOI trials (*p* = 10^−5^), as predicted, and there was no net correlation in OOO trials (*p* = 0.8), again as predicted. And in OIO trials, although the trend was not significant (p = 0.16), it was toward a positive correlation, as expected.

(3) For the activity elicited during saccades away from the RF. The model predicts that the low intensity responses evoked when the target is outside of the RF should display the same dependencies on RT as the baseline activity preceding them (Fig. 6b, d). Once again, the neural data were very similar to the model simulations (Fig. 6f, h; Supplementary Fig. 4a), and the agreement was confirmed statistically (Supplementary Fig. 4b).

In summary, the FEF activity averaged across V, VM, and M cells demonstrated varying degrees of sensitivity to RT, depending on the specific experimental condition considered, but in all cases the neuronal data conformed closely to the simulation results. Such agreement supports several key features of the model, including the competitive interactions between the target- and bias-driven responses, the limited yet visible impact of intrinsic randomness (in build-up rate) on the evoked responses, and a central hypothesis of the model — that the fluctuations in baseline at the two relevant locations are directly causal to the subsequent movement-related dynamics, and ultimately to the RTs generated.

### Heterogeneity in RT preference across FEF cells

In characterizing the functional roles of specific brain circuits, one of the main challenges is dealing with the inevitable heterogeneity of cell types and their specializations (Zeng and Sanes, 2017). Not surprisingly, single FEF neurons showed a variety of relationships to RT in the 1DR task. Remarkably, however, the model accounted for much of this diversity on the basis of a simple intuition: that the build-up rate of the target-driven activity is determined by two factors, one that is coupled to the baseline and another that is not, and those factors are weighted in various proportions at the level of single neurons. In this section we first describe the range of RT preferences measured in single FEF cells and then show that those diverse preferences naturally fall out of the elements already built into the model.

We examined the responses of individual FEF neurons during correct saccades into the RF, and found that their dependencies on RT could deviate quite substantially from those of the average population. This is illustrated with two example cells for which the maximum level of activity across trials was modulated strongly — and in opposite directions (Fig. 8a-f). To simultaneously view all the responses recorded from a given neuron, these were arranged as activity maps in which color corresponds to firing intensity and trials are ordered according to RT (Fig. 8a, d). In this way, it is clear that both cells were most active shortly before the saccade (white marks on the right) and that their firing rates were very different for the fastest versus the slowest responses (top vs. bottom trials). One cell preferred fast trials, i.e., it fired at higher rates for short RTs, whereas the other preferred slow trials, i.e., it fired at higher rates for long RTs. The contrast is also apparent when the same data are plotted as collections of firing rate traces sorted and color-coded by RT, as done in previous figures (Fig. 8b, e; compare to Fig. 6g).

For each recorded neuron, sensitivity to RT was quantified by *ρ*(*R_p_*, RT), the Spearman correlation between the peak response and RT across trials (Methods). Negative values correspond to a preference for short RTs, as for the first example cell (Fig. 8c; *ρ* = −0.58, *p* = 10^−12^), whereas positive values correspond to a preference for long RTs, as for the second example cell (Fig. 8f; *ρ* = 0.40, *p* = 10^−9^). Across our FEF sample, the resulting distribution of correlation coefficients was notable in two ways. First, it contained many more significant correlations, both positive and negative, than expected just by chance (43 of 132 cells, ^33%, were significant with *p* < 0.05, as opposed to 6.75 expected by chance; *p* =10^−22^, binomial test). Thus, a substantial fraction of the FEF neurons had robust temporal preferences, with both modulation types represented (Fig. 8g, colored points). And second, the distribution was approximately the same for all the standard FEF cell types. The proportion of positive and negative correlations, as well as the fraction of significant neurons, was statistically the same for the V, VM, M, and other categories (Fig. 8g; *p*> 0.2, binomial tests). So, as far as we could tell, the sensitivity to RT spanned a similar range for all the elements of the FEF circuitry.

**Figure 8.**
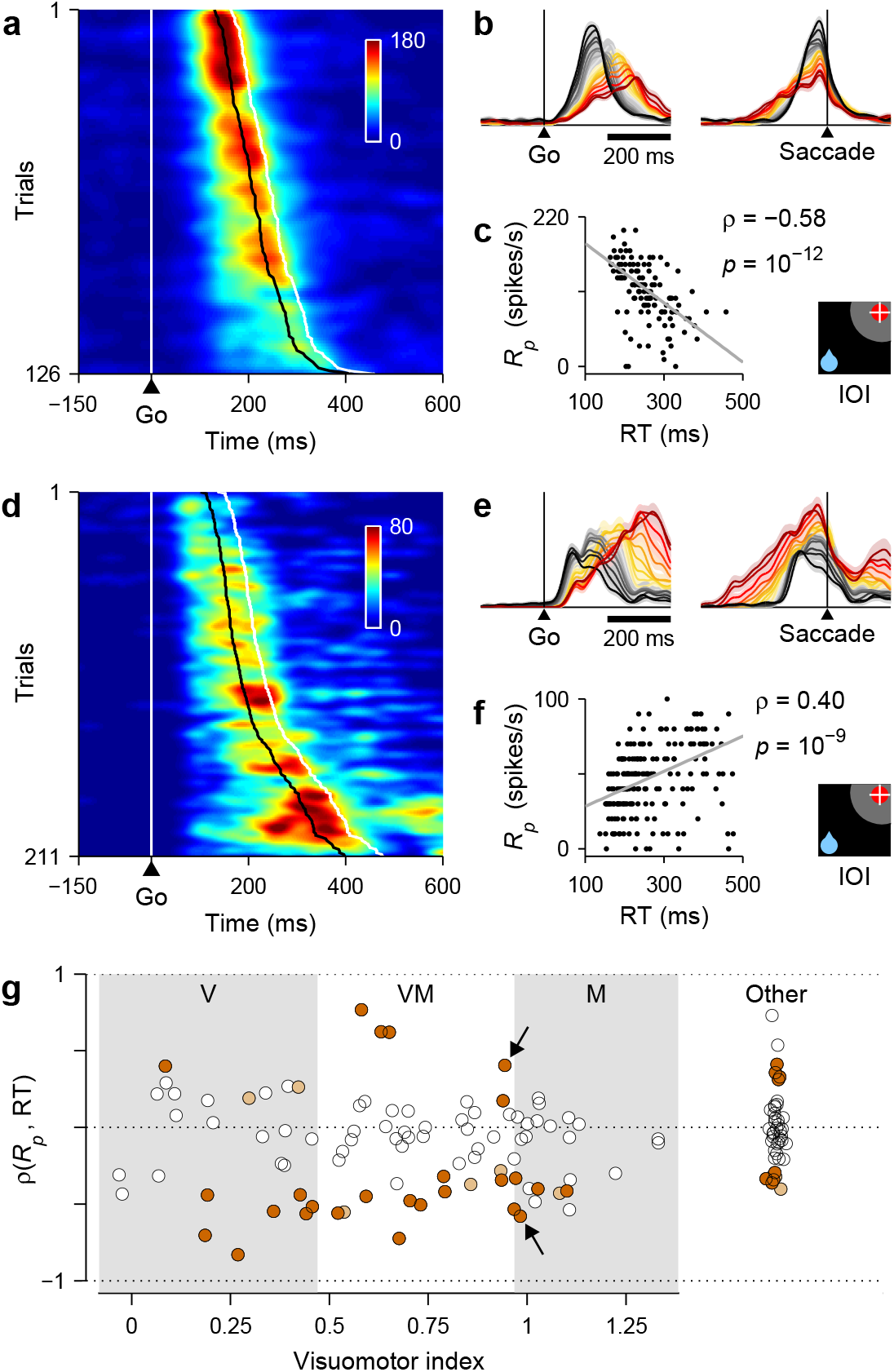
RT sensitivity of individual FEF neurons. **a-c**, Responses of a single FEF cell that fired preferentially during short RTs. All data are from IOI trials (icon). **a**, Activity map. Each row corresponds to one trial, and each trial plots firing rate (color) as a function of time (x axis), with spikes aligned to the go signal (vertical white line). Trials are ordered by RT from fastest (top) to slowest (bottom). For this cell, the time of peak activity (black marks) closely tracks saccade onset (white marks). **b**, Firing rate as a function of time, with trials sorted and color-coded by RT, as in Fig. 6. Shades indicate ±1 SE across trials. **c**, Peak response as a function of RT. Each dot corresponds to one trial. Gray line shows linear fit. Numbers indicate Spearman correlation *ρ(R_p_*, RT) and significance. **d-f**,asin**a-c**, but for a single cell that fired preferentially during long RTs. **g**, Spearman correlation between peak response and RT for all classified cells (*n* = 132). For standard cell types (V, VM, M), values on the abscissa correspond to visuomotor index; for other cells they are arbitrary. Shades demarcate ranges approximately corresponding to standard V, VM, and M categories. All correlation values are from IOI trials. Brown points indicate significant neurons (light, *p* < 0.05; dark *p* < 0.01). Arrows identify example cells in panels above.

These results explain the moderate RT sensitivity seen in the average population activity during IOI trials (Fig. 6g) as the sum of two opposing contributions from subpopulations with temporal preferences that partially offset each other. Within the fast-preferring group (*ρ* < 0), the pattern of response trajectories of many cells was qualitatively similar to that of the average population but showed more extreme variations across RT bins (Fig. 9h; compare to Fig. 6g). In contrast, only a few of the neurons within the slow-preferring group (*ρ* > 0) demonstrated a strong correlation with RT (Figs. 8d-f, 9j); for most of them, the dependence on RT, particularly during the ∼ 100 ms before movement onset, was more modest (Fig. 9i). Thus, when the responses of the fast- and slow-preferring neurons are combined, their temporal dependencies partially cancel out, and the overall population activity ends up resembling an attenuated version of the former. Similar results were obtained in the congruent condition. That is, both fast- and slow-preferring cells were also found during III trials (Fig. 9f, g), except that in that case the complementary modulations canceled out more fully upon averaging (Fig. 6e).

**Figure 9.**
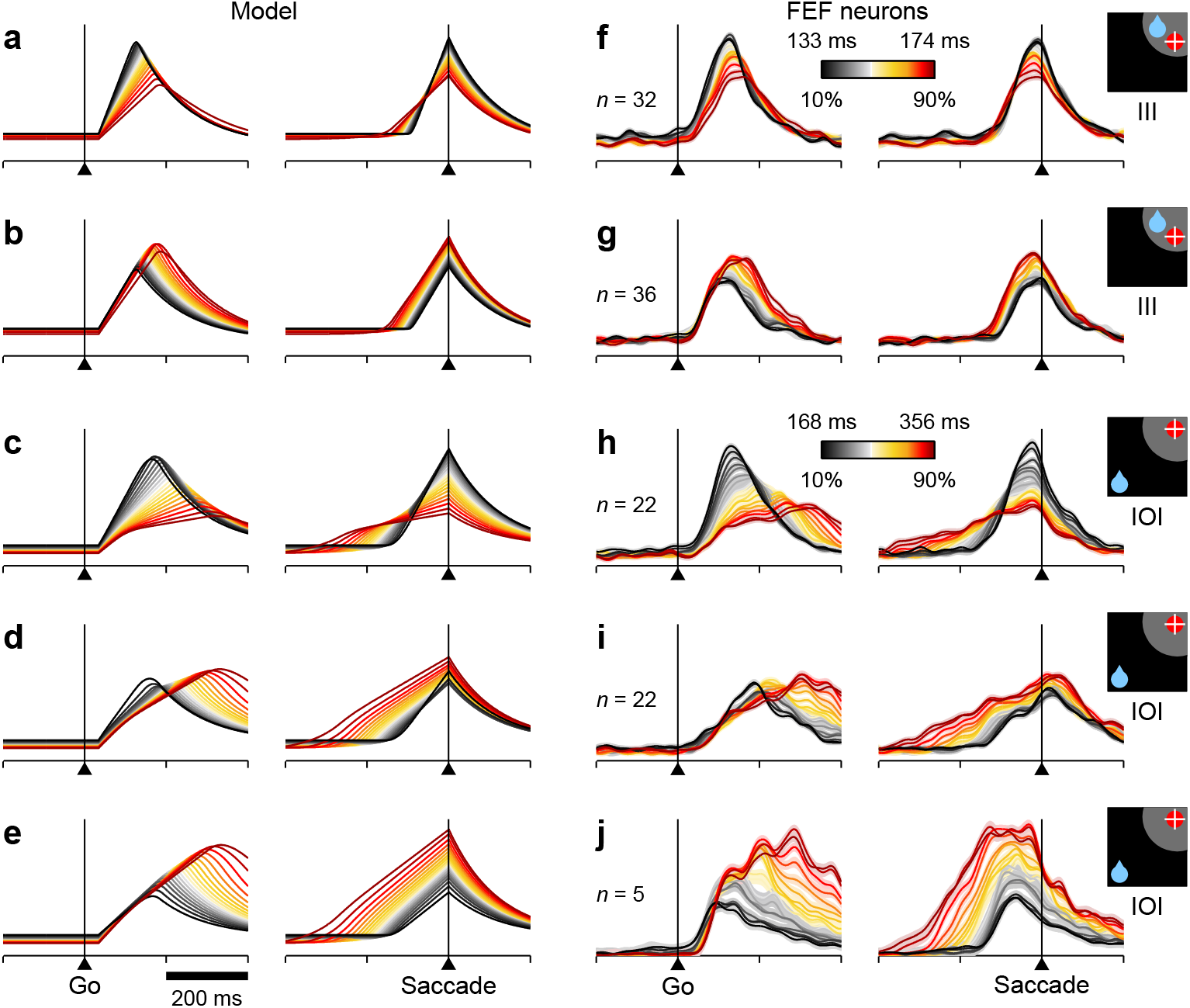
RT sensitivity in subpopulations of FEF neurons. All panels show firing rate as a function of time, with each colored curve based on a different subset of trials selected according to RT, as in Fig. 6. Only activity associated with correct saccades into the RF is displayed. a-e, Simulated activity demonstrating a clear preference for either short (**a, c**) or long RTs (**b, d, e**). The model generated fast- and slow-preferring responses in both III (**a, b**) and IOI trials (**c-e**) with varying modulation strengths. **f-j**, As in **a-e**, but based on the responses of various subsets of FEF neurons with similar RT preferences, as quantified based on *ρ*(*R_p_*, RT). Each subset is a selection from the same pool of 84 recorded neurons used in Fig. 6 (in **f**, *ρ* < 0; in **g**, *ρ* > 0; in **h**, *ρ* < 0 and *p* < 0.05; in **i**, *ρ* > 0; in **j**, *ρ* > 0 and *p* < 0.05; visuomotor index > 0.4 in all cases). Numbers of participating cells are indicated in each panel. Lighter shades behind lines indicate ± 1 SE across cells. Colorbars apply to both simulated and recorded data, either in III (colorbar in **f**) or IOI trials (colorbar in **h**).

The temporal heterogeneity just discussed was readily replicated with the model. For incongruent trials, strong modulation could be simulated favoring either short (Fig. 9c) or long RTs (Fig. 9e), but more modest temporal sensitivity like that exhibited by the majority of slow-preferring neurons could be reproduced too (Fig. 9d). In all of these cases the simulated response trajectories matched the experimental results extremely well (compare to Fig. 9h-j), both with the data aligned on the go signal, which better reflects the initial variations in build-up rate, and with the data aligned on saccade onset, which more closely tracks the activity around threshold crossing. The most visible (but still minor) discrepancy between the simulated and neuronal trajectories was due to the discontinuity of the threshold crossing event in the former, as opposed to the sharp but smooth turn around the peak of activity of the latter. Analogous results were obtained in congruent trials, for which both fast- (Fig. 9a) and slow-preferring (Fig. 9b) model responses similar to the experimental ones were also generated (compare to Fig. 9f, g). In both congruency conditions, the simulated slow-preferring responses are particularly notable because, although they are still target-driven, they would seem to require mechanisms completely different from those described earlier.

How did the model capture such wide-ranging heterogeneity? The short answer is that, without changing any of the parameters, the target-driven activity in the model, *R_T_*, can be naturally expressed as the sum of two components, like so

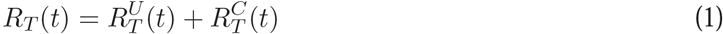

where the equality holds at every point in time. For one component (indicated by the *C* superscript), the variations in build-up rate are coupled to the fluctuations in baseline (*B_T_*), whereas for the other component (*U* superscript), the variations in build-up rate are random, uncoupled from the baseline. These two components correspond to the two neuronal types with opposite RT preferences.

To see the correspondence, consider once more the RT-sensitive responses of the FEF cells, now focusing on how the response trajectories fan out when they are aligned on the go signal: for the fast-preferring examples, the slopes of the curves increase progressively as the RTs get shorter, and the spread is visible from the moment the activity starts rising (Fig. 9c, h, left panels); in contrast, for the slow-preferring examples, all the curves start rising with approximately the same slope, and the modulation by RT begins to manifest only later, ∼100 ms after the go signal (Fig. 9d, e, i, j, left panels). According to the model, this feature, the variability of the initial build-up rate, is the fundamental mechanistic distinction between the fast- and slow-preferring FEF neurons.

A more elaborate intuition can be gleaned from the analytical expression that determines the initial build-up rate of the target-driven activity, *G_T_*, in each trial. This build-up rate can be written as

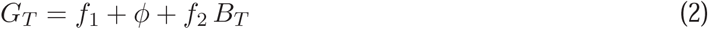

(see Equations 8, 9). Here, the terms *f*_1_, *f*_2_ and *ϕ* are not necessarily constant, but what matters is that they do not depend on the baseline at the target location, *B_T_*. The term *ϕ*, which for now is assumed to be relatively small, represents noise in *G_T_*, that is, the random fluctuations in build-up rate mentioned earlier. Intuitively, then, Equation 2 says that the initial build-up of *R_T_* is the result of two influences, a term that depends on the baseline *B_T_* plus a relatively constant drive that is independent of it. The former, *f*_2_ *B_T_*, leads to much higher variability in build-up rate across trials — and stronger covariance with RT — than the latter, *f*_1_ + *ϕ*.

Now, the coupled and uncoupled components in Equation 1 differ exclusively in their initial build-up rates, which are given by the two terms just discussed,

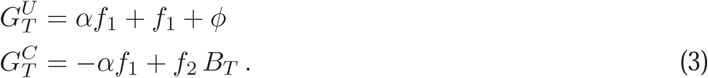

A key property of these build-up rates is that their sum, 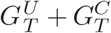, is always equal to *G_T_*, as given by Equation 2. This is true for any value of the newly introduced parameter *α*, which serves as an offset by means of which the weights of the two components may be adjusted. Splitting the target-driven activity in this way allowed us to simulate neuronal responses 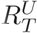 and 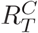 that had opposite RT preferences but were otherwise identical, as they had the same initial conditions, afferent delays, evolution equations, etc. Furthermore, by varying *α* we could vary the strength of the resulting modulation — without altering the original target-related activity, *R_T_(t)*, nor the outcomes and RTs of the competitions in any way. In other words, the split via Equations 1-3 produces paired sets of target-driven responses with different RT sensitivities, parameterized by α, but all the pairs thus generated are compatible with the same summed activity (Fig. 6) and the same distributions of outcomes and RTs (Fig. 4g-i). Based on this simple decomposition of *R* into pairs of components, the model generated a wide range of RT-sensitive responses, which were strikingly similar to those found in the FEF population.

In closing this section we emphasize the distinct role that intrinsic randomness plays in the model, and why it is necessary. During incongruent trials (or, more precisely, when *B_T_* ≤ *B_D_*) the variance in RT is so strongly coupled to the fluctuations in baseline activity that the noise in the build-up rate has a negligible impact (in Equation 2, *ϕ* ≪ *f*_2_). In that case even large variations (of ∼100%) in noise have relatively little consequence, and 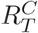, and 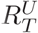 correspond almost perfectly to the fast- and slow-preferring neurons, respectively. In contrast, during congruent trials (or, more precisely, when *B_T_* > *B_D_*) a relatively large amount of independent noise (*ϕ* > *f*_2_) is necessary to reproduce the lack of sensitivity to RT in the average population activity (Fig. 6e). In that case the preferences for short and long RTs of the simulated responses (Fig. 9f, g) are noticeably sensitive to modest variations (of ∼20%) in noise, and the temporal heterogeneity of the 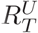 and 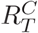 components is more complicated; the details are beyond the scope of this report. But the conclusion is clear: the large variance in RT observed during incongruent trials vastly exceeds that associated with intrinsic noise in the rising activity, and is fundamentally determined by the covert conflict between competing saccadic plans and the ensuing dynamics; whereas the much smaller variance in RT observed during congruent trials is best explained by noise that is independent of the competitive interactions, which make just a modest contribution in that case.

## Discussion

Reactive saccades are to voluntary behavior what the harmonic oscillator is to classical mechanics — the simplest non-trivial problem. And yet, the fundamental mechanisms that determine their timing have remained poorly understood. We revisited the established account of how saccades are programmed, the rise-to-threshold process, searching for any missing dynamical elements that might explain why the variance of saccadic RTs is so high even under minimalistic stimulation conditions.

We examined single-neuron activity in FEF, a cortical area whose role in saccade generation and attentional deployment is firmly established (Bruce and Goldberg, 1985; Tehovnik et al., 2000; Squire et al., 2012), and where the activity of movement-related neurons is perhaps most emblematic of the idealized rise to threshold (Hanes and Schall, 1996; Fecteau and Munoz, 2007; Stanford et al., 2010; Ding and Gold, 2012; Costello et al., 2013). We found that the three main components of this process, the baseline activity preceding target presentation, the build-up rate of the evoked response, and the saccade threshold, fluctuate in a coordinated fashion. This is already a significant departure from the simpler, standard framework in which the only source of variability (within a given experimental condition) is the build-up rate. But the problem is considerably more complicated, because by themselves, the interrelationships between these three variables are insufficient to accurately explain the variations in RT; for that, it is also critical to consider the weaker, internally-driven activity favoring saccades to alternative locations. By means of a saccadic competition model we were able to connect these and other empirical observations, which — not unlike the pieces of a puzzle — fit together just right to provide a deeper, detailed understanding of the full RT distributions in terms of circuit interactions.

Contrary to current ideas, we found that noise in the build-up rate is not the main source of RT variance. The contribution of noise was discernible but only during congruent trials, when the target-driven activity typically starts with the higher baseline and quickly advances toward threshold, with no visible competition (Fig. 3a). We concluded that, rather than intrinsic randomness, the main factor responsible for the high variance of saccadic RTs is motor conflict. In other words, the RT varies because every saccade is the result of a motor selection process.

During reactive saccades, it may seem as if only one motor plan is possible, but this is rather illusory. When the target appears, oculomotor activity starts growing in response to it, but this ramping process (represented by *R_T_* in the model) does not start from the same neutral state every time; instead, it occurs while other incipient, internally-driven motor plans (represented by *R_D_*) are also developing, and the time necessary to resolve the ensuing conflict depends on how advanced and how congruent those budding, bias-driven plans are relative to the target-driven response. Making the target unique, highly visible, and task-relevant minimizes potential variance related to the sensory detection step and enhances the priority of the target-related plan, but still leaves those alternative internal plans relatively unconstrained. What the 1DR task does is align the internal bias with a specific direction — that of the expected reward — and it is under those conditions that the motor selection process becomes more apparent (Fig. 3). The model revealed completely novel mechanistic details of this motor competition but, on average, its manifestations during the rise to threshold were nevertheless quite subtle, particularly during correct saccades.

Consider, for example, how the longest RTs in correct trials are produced (Fig. 4b, h). That was quite a puzzle. In that case the target-driven activity, which starts with a lower baseline than the opposing plan, must somehow rise at about the lowest possible build-up rate and yet still win the competition by a large margin. The solution is for the target-driven response to rise quickly initially, suppress the opposing plan early on, and then slow down immediately afterward — all of which happens when the competing plans are far from threshold. This is a major departure from standard choice models, in which the outcome is not determined until activity is close to or at threshold. It means that critical dynamical interactions may occur at quite low levels of activity, where they are less readily apparent and much more difficult to characterize without prior knowledge of their signature features.

The model has numerous moving parts, but consider the scope of the data that it reconciled (Methods, *Correspondence between data and model parameters*). Behaviorally, the model generated errors at the appropriate rates and reproduced three distinct RT distributions in their entirety (Fig. 4g-i); and neurophysiologically, it replicated the average response trajectories in all conditions (Fig. 4d-f), the isolated effect of the spatial bias (Fig. 5c-f), the dependence on RT of the average activity (Fig. 6), and most remarkably, the responses of individual FEF neurons, which showed a wide range of RT preferences (Fig. 9). Thus, a large number of disparate observations are subsumed into one coherent, compact framework for resolving the kind of saccadic conflict that must typify naturally occurring oculomotor behaviors.

### The threshold varies — but not as we thought

According to the model, the saccadic threshold is not constant, but rather fluctuates quite dramatically. Such fluctuations are visible when comparing average movement-related activity across experimental conditions (Edelman and Goldberg, 2001; Thompson et al., 2005; Heitz and Schall, 2012; Jantz et al., 2013; Jagadisan and Gandhi, 2017) (Figs. 3, 5). However, characterizing them in individual trials is extremely difficult when neurons are recorded one at a time, which is why the model is critical. Our results indicate that the threshold fluctuations are not random, but rather are coordinated with other elements of the circuitry, and although the proposed dynamics remain to be directly tested in other saccadic tasks, substantial agreement can already be found with extant data. For instance, movement-related activity preceding memory-guided saccades or anti-saccades is considerably weaker than for stimulus-driven saccades (Edelman and Goldberg, 2001), consistent with a lower threshold for internally-driven motor plans, and the same is true for saccades that are triggered by blinks (Jagadisan and Gandhi, 2017). The presaccadic activity measured during visual search is less vigorous for incorrect than for correct responses to the same location (Thompson et al., 2005), presumably because the former involve a stronger internal (and erroneous) drive that promotes a lower threshold. And, again in the context of visual search, one study (Heitz and Schall, 2012) showed multiple differences in FEF activity across task conditions similar to those found here; that is, when the mean RT was shorter, the baseline, build-up rate, and threshold were all higher.

These findings place significant constraints on the trigger mechanism that converts a saccadic plan into a committed, uncancelable command (Hanes and Schall, 1996; Lo and Wang, 2006; Brown et al., 2008). For instance, a popular idea is that adjustments in threshold may serve to trade speed against accuracy during choices (Lo and Wang, 2006; Bogacz et al., 2010; Heitz and Schall, 2012; but see Salinas et al., 2014; Thura et al., 2014; Thura and Cisek, 2016, 2017). This is simply because, everything else being equal, activity that ramps toward a higher threshold should take longer to reach it, thus providing more time for deliberation. By exactly the same logic, it has also been suggested that control of the baseline could serve the same purpose (Reddi and Carpenter, 2000; van Veen et al., 2008; Bogacz et al., 2010). However, the results in the 1DR task are entirely antithetical to these notions: congruent trials produce shorter RTs and higher accuracy than incongruent ones, largely because of strong modulation of the build-up rates and, if anything, in spite of a longer excursion from baseline to threshold (Figs. 3, 4). This is much more in line with theoretical studies (Salinas and Abbott, 1996; Chance and Abbott, 2002; Salinas, 2003; York and van Rossum, 2009) showing that, in general, the background activity in recurrent circuits is likely to have profound dynamical and amplification effects on evoked responses. Large variations in baseline and threshold do occur in the 1DR task, but they are linked to each other and to variations in build-up rate in ways that are inconsistent with the standard characterization of the speed-accuracy tradeoff.

These considerations are significant because, although the threshold is a universal feature of decision-making models, the strongest evidence of its existence is precisely the behavior of oculomotor neurons in FEF and SC (Hanes and Schall, 1996; Lo and Wang, 2006; Brown et al., 2008; Stanford et al., 2010; Ding and Gold, 2012). In other, related circuits either no threshold is apparent (Ding and Gold, 2010; Stuphorn et al., 2010) or its implementation is much less evident (Afshar et al., 2011;Hayden et al., 2011).

### It is all about the base

Sumner (2011) has pinpointed why explaining saccadic latency distributions is so challenging: it is well established that Gaussian variability in the build-up rate of a rise-to-threshold process accurately reproduces their characteristic skewed shapes (Carpenter and Williams, 1995; Hanes and Schall, 1996; Fecteau and Munoz, 2007), but most factors known to affect saccadic RT in simple tasks are normally modeled as baseline shifts (Sumner, 2011). Our results suggest that the dichotomy is false. The fluctuations in baseline, build-up rate, and RT are inextricably linked (Figs. 5, 6, 7), and there is no simple formula for going from one of these variables to another; the transformations are nonlinear, involve alternate, covert motor plans, and depend on multiple neural mechanisms acting in concert.

Notably, according to the model, the major source of randomness across trials is the variability of the baselines (Equation 5). Aside from the target-driven build-up rate, which is subject to additional, independent fluctuations (*ϕ* in Equation 2), all quantities are either constant or deterministically related to the baselines. This is a simplification, of course. It is certainly possible to add independent noise to other components of the model (e.g., the afferent delays) without substantially altering the results; what is remarkable, though, is that this is not necessary. In the incongruent condition, in particular, nearly all of the variance observed experimentally — in RT, saccadic choice, threshold level, and in the build-up rates and peak responses of the neurons — results from the computational amplification of the initial fluctuations in baseline. Arguably, the baselines reflect multiple cognitive elements, including expectation, anticipation, and the allocation of attention and other resources (Bruce and Goldberg, 1985; Coe et al., 2002; Maunsell, 2004; Rao et al., 2012; Zhang et al., 2014; Thura and Cisek, 2016), in agreement with the effects of sub-threshold microstimulation (Glimcher and Sparks, 1993; Dorris et al., 2007; Squire et al., 2012). In the model, they set the initial spatial priorities, and thereafter the circuit dynamically blends them with the incoming sensory signal to produce the next saccade. In this formulation, the resulting variance in RT is not the consequence of noisy representations or sloppy computations, but rather the signature of a well-oiled motor-selection machine (Najemnik and Geisler, 2005; Tian et al., 2016).

The model proposes a tight relationship between initial state and subsequent oculomotor dynamics, and interestingly, mounting evidence demonstrates a similar phenomenon in motor cortex whereby the initial neural state is predictive of an ensuing arm movement and of the trajectories of the underlying neural signals over time (Churchland et al., 2006, 2010, 2012; Afshar et al., 2011; Ames et al., 2014; Sheahan et al., 2016; Stavisky et al., 2017). In that case the dynamics have a strong oscillatory component (Churchland et al., 2012) and develop within a very high-dimensional space, such that the preparatory activity is only weakly related to specific kinematic parameters (Churchland et al., 2006, 2010). Saccades are simpler because they are lower dimensional and largely ballistic, and because the activity of any given cell generally corresponds to a fixed movement vector. However, we propose that their dynamical behavior is qualitatively similar in that the initial state of the system — i.e., the configuration of baseline levels across RFs during fixation — fundamentally determines its subsequent temporal evolution, including its interaction with new incoming sensory information (Sheahan et al., 2016) and the eventual outcome (Churchland et al., 2006; Afshar et al., 2011).

Overall, the current results suggest that, when oculomotor circuits receive new visual information, ongoing saccadic plans and internal settings (e.g., threshold level, attentional locus) radically shape the impact of that information, even when it is behaviorally relevant, expected, and unambiguous. Deeper understanding of the underlying network dynamics will be critical for further elucidating how saccades are triggered and, more generally, how perceptually-guided choices are made.

## Methods

### Subjects and setup

Experimental subjects were two adult male rhesus monkeys (*Macaca mulatta*). All experimental procedures were conducted in accordance with the NIH Guide for the Care and Use of Laboratory Animals, USDA regulations, and the policies set forth by the Institutional Animal Care and Use Committee (IACUC) of Wake Forest School of Medicine.

An MRI-compatible post served to stabilize the head during behavioral training and recording sessions. Analog eye position signals were collected via scleral search coil (Riverbend Electronics) and infrared tracking (EyeLink 1000, SR Research). Stimulus presentation, reward delivery, and data acquisition were controlled by a purpose designed software/hardware package (Ryklin Software). Target stimuli were displayed via a 48x42 array of tri-color light-emitting diodes. Saccade onset was identified as the time at which eye velocity exceeded 50°/s; having detected the start of a saccade, its end was identified as the time at which eye velocity fell below 40°/s. Eye movements were scored as correct if the saccade endpoint fell within 5° of the target stimulus.

Neural activity was recorded using single tungsten microelectrodes (FHC, 2-4 MΩ) driven by a hydraulic microdrive (FHC). Individual neurons were isolated based on the amplitude and/or waveform characteristics of the recorded and filtered signals (FHC; Plexon, Inc). Putative FEF neurons were selected from areas in which saccade-like movements could be evoked by low current microstimulation (70 ms stimulus trains at 350 Hz, with amplitude equal to 50 *μ*A). (Bruce and Goldberg, 1985; Bruce et al., 1985; Costello et al., 2013). Neurons were recorded unilaterally in both monkeys.

### Behavioral tasks

In the 1DR task (Fig. 1a), all trials began with the appearance of a centrally located stimulus. Monkeys had to maintain their gaze on this fixation spot for 1000 ms. The disappearance of the fixation spot (go signal) was simultaneous with the appearance of a second, target stimulus in one of four possible positions (up, down, left, right in Fig.1a), which were chosen based on the RF of each recorded neuron. Subjects were required to make a saccade to the peripheral target within 500 ms of the go signal in order to receive a liquid reward. Target locations varied pseudorandomly from trial to trial. In each block of trials, only one of the four target locations was associated with a large reward; the other three were unrewarded (Monkey G), or yielded a much smaller reward (Monkey K). For brevity we refer to these simply as the “rewarded” and “unrewarded” locations. The rewarded location changed pseudorandomly from one block to another. Block length was highly variable (range: 10-140 trials); the average was 70 trials per block.

In the all-directions-rewarded task (ADR), the events were the same as in the 1DR, but the four target locations were rewarded equally (Fig. 1b). Blocks of ADR trials were run sporadically, interleaved with those of 1DR trials.

In the delayed-saccade task, each trial began with fixation of a central spot, followed by the appearance of a single stimulus at a peripheral location during continued fixation. After a variable delay (500, 750, or 1000 ms), the fixation spot was extinguished (go signal) and the subject received a liquid reward if a saccade was made to the peripheral target. In each experimental session, the delayed-saccade task was run first to locate the RF of the recorded neuron, and subjects performed the 1DR task after the initial spatial characterization.

The RT was always measured from the go signal until the onset of the saccade.

### Trial selection

For all analyses not specifically examining sequential effects and block transitions, we discarded the first 8 trials of each 1DR block, during which the monkeys may have been discovering the new rewarded location (Supplementary Fig. 1a). This guaranteed that all behavioral and neural metrics reflected a stable expectation, and that erroneous saccades were not due to spatial uncertainty. More stringent exclusion criteria produced qualitatively similar results.

### Continuous firing activity

Continuous (or instantaneous) firing rate traces, also known as spike density functions, were computed by convolving evoked spike trains with a Gaussian function (σ = 15 ms) with unit area. Continuous mean traces for each neuron were generated by averaging across trials. To produce equivalent population responses (e.g., Fig. 3a-c), the continuous traces of individual cells were first normalized by each neuron's overall maximum firing rate and were then averaged across neurons.

To visualize how RT modulated the activity of each cell, families of firing rate traces ordered and color-coded by RT were generated (Fig. 8b, e). For this, the trials in the relevant experimental condition (e.g., IOI, III) were sorted by RT and distributed over 20 evenly-spaced, overlapping bins defined by RT quantiles, where each bin contained 20% of the trials. Thus, the first bin was centered on the 10th percentile and included the fastest 20% of the recorded trials, the next bin included the 20% of the trials around the 14th percentile, and so on, with the last bin centered on the 90th percentile and including the slowest 20% of the recorded trials. Then, a continuous firing rate trace was produced for each of the 20 RT bins/quantiles. To generate equivalent families of curves for populations of neurons (Fig. 6f-j), the traces for each participating cell were first normalized by that cell's overall maximum firing rate, and then, for each quantile, a population trace was compiled by averaging across neurons. This method, as opposed to standard binning using fixed RT values, reveals more clearly the actual modulation range of the neural responses and their smooth dependence on RT, because the numbers of trials and neurons remain nearly constant across bins. Also, if the dependence on RT is monotonic, as the data indicate, this procedure can only *under*estimate the magnitude of the modulation. For the simulated data, families of response curves ordered by RT were generated in the same way.

For each neuron, an activity map (Fig. 8a, d) for a given condition (e.g., IOI, III) was assembled by aligning all spike trains to the go signal, converting each one to a continuous firing rate, sorting the trials by RT, putting the sorted firing rate traces into a single matrix, and displaying the matrix as a heat map with color indicating intensity. For display purposes, activity maps were also smoothed with a Gaussian function in the vertical direction, i.e., across trials (*σ* = 2 trials), but this was exclusively for ease of viewing; trials were kept independent in all analyses.

### Peak response

The peak response in each trial, *R_p_*, is well suited for quantifying how responsivity changes with RT because it is insensitive to the alignment of the spike trains. This is in contrast to the standard firing rate calculated in a fixed time window anchored to a particular task event (e.g., go or saccade onset). With an anchored window it is possible to observe a spurious dependence on RT simply because of temporal misalignment across trials — but not so with the peak response.

For each neuron, the value of *R_p_* in each trial was equal to the cell's firing rate computed in an interval centered on the time point *T_p_*, which we call the time of peak response. This is simply the time along a trial (with the go signal at *t* = 0) at which the cell was most likely to fire at the highest rate (Fig. 8a, d, black marks). *T_p_* is described as a linear function of RT.

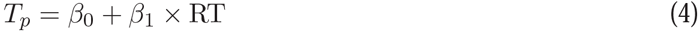

where the coefficients *β*_0_ and *β*_1_ characterize the timing of each neuron. For example, for a typical visual cell with *β*_0_ = 80 and *β*_1_ =0, the maximum rate in a trial is observed 80 ms after the go signal, regardless of RT, whereas for a typical movement-related cell with *β*_0_ = −30 and *β*_1_ = 1, the highest discharge occurs 30 ms before the saccade.

The coefficients *β*_0_ and *β*_1_ were obtained in two steps: (1) finding, from the activity map of the cell, the maximum instantaneous firing rate in each trial and the time, relative to the go signal, at which that rate was achieved, *T*_max_, and (2) fitting *T*_max_ as a linear function of RT. All trials in which a saccade was made into the cell's RF were included, regardless of bias condition. The coefficients resulting from the fit, i.e., the intercept and slope, were *β*_0_ and *β*_1_. Finally, to determine *R_p_* in a given trial, first, the corresponding *T_p_* was found by plugging the RT from that trial into Equation 4; then a firing rate was calculated by counting the spikes in a time window centered on *T_p_* and dividing by the window length. The result was *R_p_*. The window length was 100 ms for most cells (∼80%) but was set to 200 ms for a minority that had more prolonged responses (e.g., postsaccadic cells).

### Statistical analyses

All data analyses were performed using customized scripts in MATLAB (The MathWorks, Natick, MA). For comparisons across any two conditions, significance (*p* < 0.05) was typically evaluated via permutation tests for paired or unpaired samples (Siegel and Castellan, 1988), as appropriate. Because 100,000 permutations were used, the smallest significance value in this case is reported as *p* < 10^−5^.

The relationship between neuronal activity and RT for each neuron was evaluated separately for each experimental condition (e.g., III, IOI) using the Spearman rank correlation coefficient. The Matlab function corr was used for this. This coefficient serves to identify any monotonic relationship between two variables. As measures of activity we considered, for each cell, the baseline response, *R_b_* (firing rate in a 250 ms window preceding the go signal); the mean response, *R_m_* (firing rate computed over the full RT interval); the presaccadic response, *R_sac_* (firing rate in a 50 ms window preceding saccade onset); and the peak response, *R_p_* (described above). The correlation between *R_p_* and RT is denoted as *ρ*(*R_p_*, RT), and similarly for other activity measures. Neurons with *ρ*(*R_p_*, RT) < 0 and *ρ*(*R_p_*, RT) > 0 were designated as fast- and slow-preferring, respectively.

### Neuronal classification and visuomotor index

The 132 FEF neurons were classified by comparing their responses (mean firing rate in windows of 100 – 250 ms) during fixation, during the RT interval, and after the saccade. Multiple-comparison tests were performed via ANOVA. Accordingly, cells that were maximally active before the go signal were classified as fixation neurons (*n* = 12); cells that responded significantly above baseline, but only after the saccade, were deemed postsaccadic (*n* = 18); and neurons that started responding shortly after the go signal and that were still significantly active after the saccade were deemed wide-profile (*n* = 10). The latter group could conceivably have been included in the visuomotor category described below, but given their peculiar lack of sensitivity to the saccade, they were analyzed separately.

The rest of the neurons (*n* = 92) had standard visuomotor properties and responded significantly above baseline between the go signal and saccade onset. A visuomotor index, which was equated to the coefficient *β*_1_ in Equation 4, was used to characterize them (Fig. 8g). This coefficient naturally serves as a visuomotor index because it describes the degree to which a neuron is activated by a stimulus in its RF (in which case *β*_1_ ≈ 0) versus by an eye movement toward it (in which case *β*_1_ ≈ 1) — which is the classic criterion used to classify FEF cells (Bruce and Goldberg, 1985). So, based on their *β*_1_ values, the remaining 92 neurons were classified as either visual (V; *n* = 26), visuomotor (VM; *n* = 43), or motor (M; *n* = 23). As expected from previous studies (Bruce and Goldberg, 1985; Costello et al., 2013), during delayed-saccade trials, the neurons thus classified as V responded briskly to the presentation of the target stimulus in the RF and gradually decreased their activity thereafter; M neurons showed no activity linked to stimulus presentation but responded intensely just before movement onset; and the cells in the VM group showed both visual and presaccadic activation.

### RT matching

To tease apart the effects of RT and spatial bias on FEF activity, we devised a procedure for equalizing the RT distributions of the congruent and incongruent conditions. For each recorded cell, the observed distributions in III and IOI trials (Fig. 5a) were sub-sampled as follows. An III trial was selected randomly and the IOI trial with the most similar RT was identified; then, if the RT difference was smaller than 15 ms, the two trials were accepted into the respective sub-samples and removed from the original pools, or else the III trial was discarded. After probing all the III trials like this, the resulting sub-sampled pools (Fig. 5b) contained equal numbers of trials with virtually identical RT sets. To account for variations due to random resampling, all results based on RT matching were repeated 50 times and averaged.

### Saccadic competition model

The model consists of two populations of FEF neurons that trigger saccades toward locations *T* (where the target stimulus appears) and *D* (diametrically opposite to *T*), their activities represented by variables *R_T_* and *R_D_*. After the go signal is given, both motor plans start increasing, and the first one to reach a threshold Θ wins the competition, thus determining the direction of the evoked saccade and the RT. If *R_T_* wins, the saccade is correct, toward *T*, whereas if *R_D_* wins, the saccade is incorrect, toward *D*. Each simulated race corresponds to one trial of the 1DR task. The actual trajectories followed by *R_T_* and *R_D_* in each trial are dictated by the dynamics and interactions described below.

In each simulated trial, three key quantities need to be specified before the race between *R_T_* and *R_D_* can take place: the baseline firing levels, which serve as the initial values for *R_T_* and *R_D_*, the initial build-up rates of the two motor plans, and the threshold Θ. Simulated neural responses are scaled so that the firing activity at threshold is around 1. The baselines for the target and distracter locations, *B_T_* and *B_D_*, are specified first, drawn according to the following expressions,

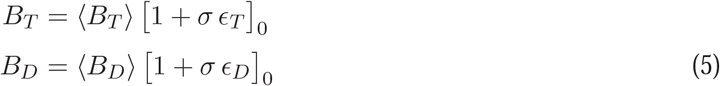

where the variability across trials is determined by *σ* = 0.28, and *∊_T_* and *∊_D_* are random Gaussian samples (different for each trial) with negative correlation equal to −0.5, zero mean, and unit variance. Here and in the expressions below, the square brackets with a subscript indicate that there is a minimum floor value beyond which the argument cannot drop, that is, [*x*]_*a*_ = max{*x, a*}. This ensures, for instance, that quantities commensurate with firing activity are not negative. The mean baseline levels in Equation 5, 〈*B_T_*〉 and 〈*B_D_*〉, are set to be consistent with the spatial bias: in incongruent trials 〈*B_T_*〉 = 0.16 and 〈*B_D_*〉 = 0.34, so the target side has the lower baseline on average, whereas in congruent trials 〈*B_T_*〉 = 0.34 and 〈*B_D_*〉 = 0.16, so the target side takes the higher value. Stated differently, the higher mean baseline is always assigned to the rewarded location.

Once the baselines are drawn, the other key quantities, the threshold and the initial build-up rates, can be set for the trial. The threshold is given by

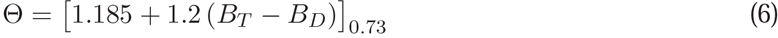

where Θ cannot drop below 0.73, as indicated by the square brackets. This expression means that the threshold for triggering a saccade increases with the baseline level on the target side and decreases with the baseline level on the opposite side, but cannot be less than a certain minimum.

The initial build-up rates (or gains) of the motor plans also depend on the baselines. For the internally-driven plan, *R_D_*, the build-up rate is

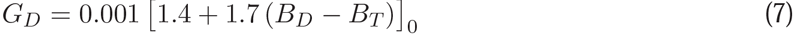

so the activity favoring *D* rises most steeply when the baseline at that location is high and the baseline at the target location is low. For the target-driven motor plan, *R_T_*, the rise in activity has two distinct regimes. In the first one, when *B_T_* ≥ *B_D_*, the initial build-up rate is given by

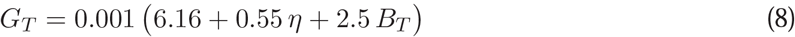

where *η* is a random Gaussian sample (zero mean, unit variance) that varies stochastically across trials and represents intrinsic, baseline-independent fluctuations in build-up rate. In the second regime, when *B_T_* < *B_D_*,

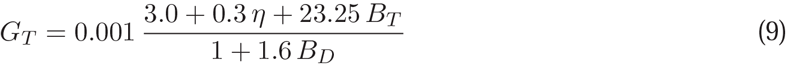

where the numerator has the same form as in Equation 8 but now *B_D_* appears in the denominator. The rationale for using two distinct expressions for *G_T_* is simply that the rise of the target-driven activity is very different when *R_T_* already is above the competition before the race begins versus when it starts below the competition (Fig. 3a, b). In the former case (*B_T_* ≥ *B_D_*, regime 1) the rise is always steep, the dependence on *B_T_* is weak, and there is no further opposition from the *D* plan (note the absence of a *B_D_* term in Equation 8). In contrast, when the target side starts at a disadvantage (*B_T_* < *B_D_*, regime 2) the initial build-up rate depends strongly on the actual baseline level, *B_T_*, and there is sizable competition from the *D* plan, instantiated as divisive suppression (note the dependence on *B_D_* in the denominator of Equation 9).

It is important to realize that, because the baselines fluctuate across trials (Equation 5), in general, Equation 8 applies most often, but not uniquely, to congruent trials; and similarly, Equation 9 applies most often, but not uniquely, to incongruent. The build-up rate *G_T_* depends only on the baseline values themselves, regardless of the label assigned to the spatial configuration of each trial. In other words, the local competition process has no knowledge of what determines the baselines, it simply takes them as input and evolves accordingly.

The main variables, *R_T_* and *R_D_*, which represent the activities of the competing populations, are updated in each time step Δ*t* (set to 1 ms) as follows

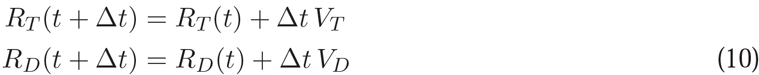

where *V_T_* and *V_D_* are the instantaneous build-up rates. Initially, i.e., during fixation, the activities are equal to their respective baseline values, and they remain constant until the target/go signal is presented (at *t* = 0). Thus, the initial conditions are *R_T_* = *B_T_*, *R_D_* = *B_D_*, *V_T_* = 0 and *V_D_* = 0. The motor plans start advancing thereafter, but not right away, because there is an afferent delay between the go signal and the actual onset of ramping activity. The target-driven plan, *R_T_*, starts rising after a short delay *A_T_* = 35 ms, and does so with the build-up rate prescribed by Equation 8 or 9, whichever applies (which can be coded as: if *t* ≥ *A_T_*, then *V_T_* = *G_t_*). The bias-driven plan, *R_D_*, starts rising after a slightly longer delay *A_D_* = 50 ms, but at the beginning it is partially inhibited by the cue presentation. During this partial inhibition, which occurs between *I*_on_ = 40 and *I*_off_ = 155 ms, *R_D_* rises slowly, at 38% of its nominal build-up rate, *G_d_* (if *t* ≥ *A_D_* and *t* ∈ [*I*_on_, *I*_off_], then *V_D_* = 0.38 *G_D_*). This dynamic is based on evidence indicating that the abrupt appearance of a visual stimulus, the target in this case, briefly interrupts or suppresses ongoing saccadic plans (Reingold and Stampe, 2002; Dorris et al., 2007; Stanford et al., 2010; Bompas and Sumner, 2011; Buonocore and McIntosh, 2012; Buonocore et al., 2017), a phenomenon known as “saccadic inhibition” or the “remote distracter effect”. After the inhibition period has elapsed, the bias-driven activity may rise in full force (if *t* > *I_off_*, then *V_D_* = *G_D_*).

The two motor plans then keep advancing until one of them reaches threshold. However, their build-up rates may change mid-flight as one plan goes past the other. These changes are dictated by two rules that describe the two possible ways in which the competition may end.

<u>Rule 1</u> (*T* wins): if the target-driven firing rate, *R_T_*, exceeds the competing one at any point after its afferent delay has elapsed, then two things happen. First, *R_D_* is fully suppressed, so it stops increasing altogether (if *t* > *A_T_* and *R_T_* > *R_D_*, then *V_D_* = 0). And second, the build-up rate of the *T* plan is adjusted so that

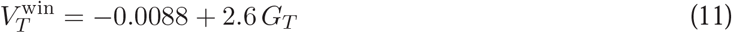

(if *t* > *A_T_* and *R_T_* > *R_D_*, then 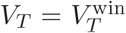). In this case, *R_T_* wins the race and the evoked saccade is correct, toward the target. The coefficients in Equation 11 are such that 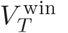 is typically smaller than *G_T_*. This means that the target-driven motor plan typically slows down after it overtakes the competition, and the lower its initial build-up rate, the more it slows down. This change in build-up rate represents the difficulty, or cost, of resolving the conflict for the *T* plan.

<u>Rule 2</u> (*D* wins): if the internally-driven firing rate, *R_D_*, exceeds the competing one at any point after its afferent delay has elapsed and outside of the transient inhibition interval, then *R* can no longer advance past *R_D_*. In this case *R_D_* simply continues to rise, winning the race without any further change in its build-up rate. The evoked saccade is incorrect, away from the target. The target-driven motor plan also keeps rising, but may suffer a minimal amount of suppression (the amount needed to ensure that *R_T_* stays below *R_D_* for the remainder of the trial).

Finally, to account for the characteristic fall in activity seen postsaccadically in FEF, after the winner reaches threshold both motor plans decay exponentially toward a firing level of 0.2 with a time constant of 120 ms.

Note that the model generates different outcomes and RTs based only on three quantities (random numbers) that vary across trials: *∊_T_* and *∊_D_*, which determine the baseline firing levels (Equations 5), and *η*, which adds independent noise to the build-up rate of *R_T_* (Equations 8, 9). There are no other sources of variability.

### Correspondence between data and model parameters

All parameter values in the model were adjusted to fit the experimental data. Each parameter typically has multiple effects, often on both the behavioral and neurophysiological responses simulated. For example, the afferent delays determine the short pause between the go signal and the onset of the rise to threshold (seen experimentally in Fig. 3a-c, left panels), but they also pin the left tails of the RT distributions (Fig. 4g-i) because they determine the shortest possible RTs. Similarly, the variance of the baselines (*σ*) generally determines the correlation between activity and RT (Figs. 6, 3), but it also influences the widths of the RT distributions during incongruent trials (Fig. 4h, i).

With this in mind, note that the parameters in Equations 5, 6 were primarily set to match the baseline and threshold values measured from the FEF population across conditions (Supplementary Fig. 3). The parameters in Equations 7, 8, and 9 mainly determine the shapes of the RT distributions for incorrect incongruent, correct congruent, and correct incongruent trials, respectively (Fig. 4g-i). The parameters that describe the stimulus-driven suppression of *R_D_* determine the frequency of incorrect saccades, the shape of the corresponding RT distribution, and the shape of the simulated response trajectories in those incorrect trials. Finally, the parameters associated with Rule 1 scale the RT distribution for correct incongruent trials, and also determine the steepness of the simulated target-driven rise in activity.

In all, 22 model parameters were adjusted to satisfy 23 basic experimental constraints: 6 baseline and 6 threshold values across conditions (Supplementary Fig. 3), 2 error rates, and 3 RT distributions (Fig. 4g-i), each of which requires, at a minimum, 3 parameters to be characterized. But note that the model accounted for many more features (i.e., degrees of freedom) in the data, pertaining to the specific shapes of response trajectories, the effect of RT equalization, the correlation between firing activity and RT before and after target onset, and the response heterogeneity across individual FEF neurons.

---

**Supplementary Figure 1.**
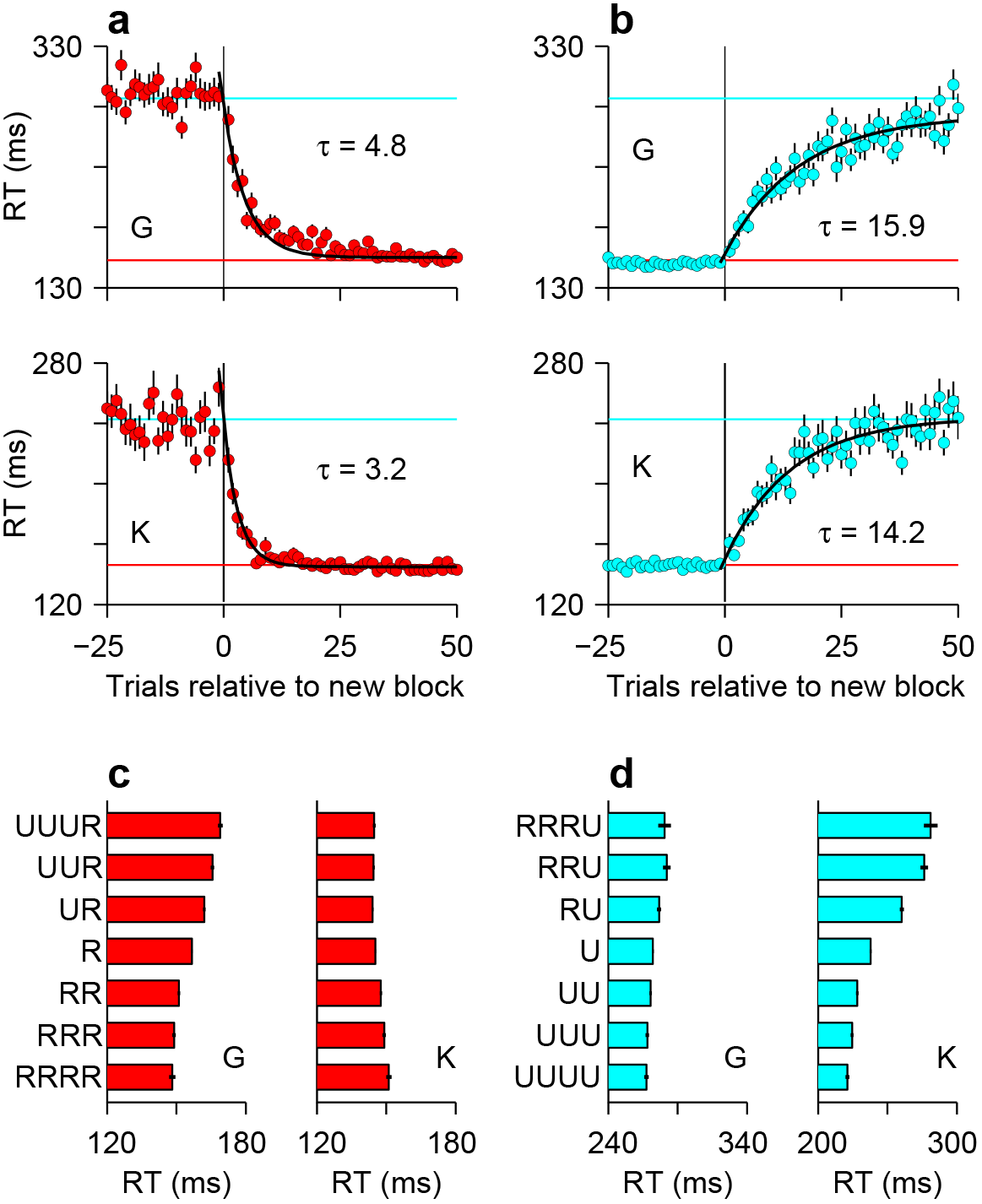
Variations in spatial bias over time. Analyses of the sequential responses made by the two animals confirmed that they were strongly aware of which location was the rewarded one. **a, b**, Time course of bias adaptation after a change in rewarded location. Mean RT (± 1 SE) for saccades to a fixed location as a function of the number of trials before or after a change in reward status (i.e., block transition). Transitions from unrewarded to rewarded (**a**, red) are faster than from rewarded to unrewarded (**b**, cyan). Horizontal lines indicate asymptotic RT values. Black lines are exponential fits with time constants as indicated (*τ*; units are trials). **c, d**, Trial history effects. Mean RTs (± 1 SE) to rewarded (**c**, red) and unrewarded (**d**, cyan) targets are shown conditioned on various preceding sequences of rewarded (R) and unrewarded (U) trials. Note systematic changes in RT, which are somewhat idiosyncratic to each animal, as each condition is repeated. For calculating these trial-history effects, the first 15 trials after each block transition were excluded. All data are from correct responses.

**Supplementary Figure 2.**
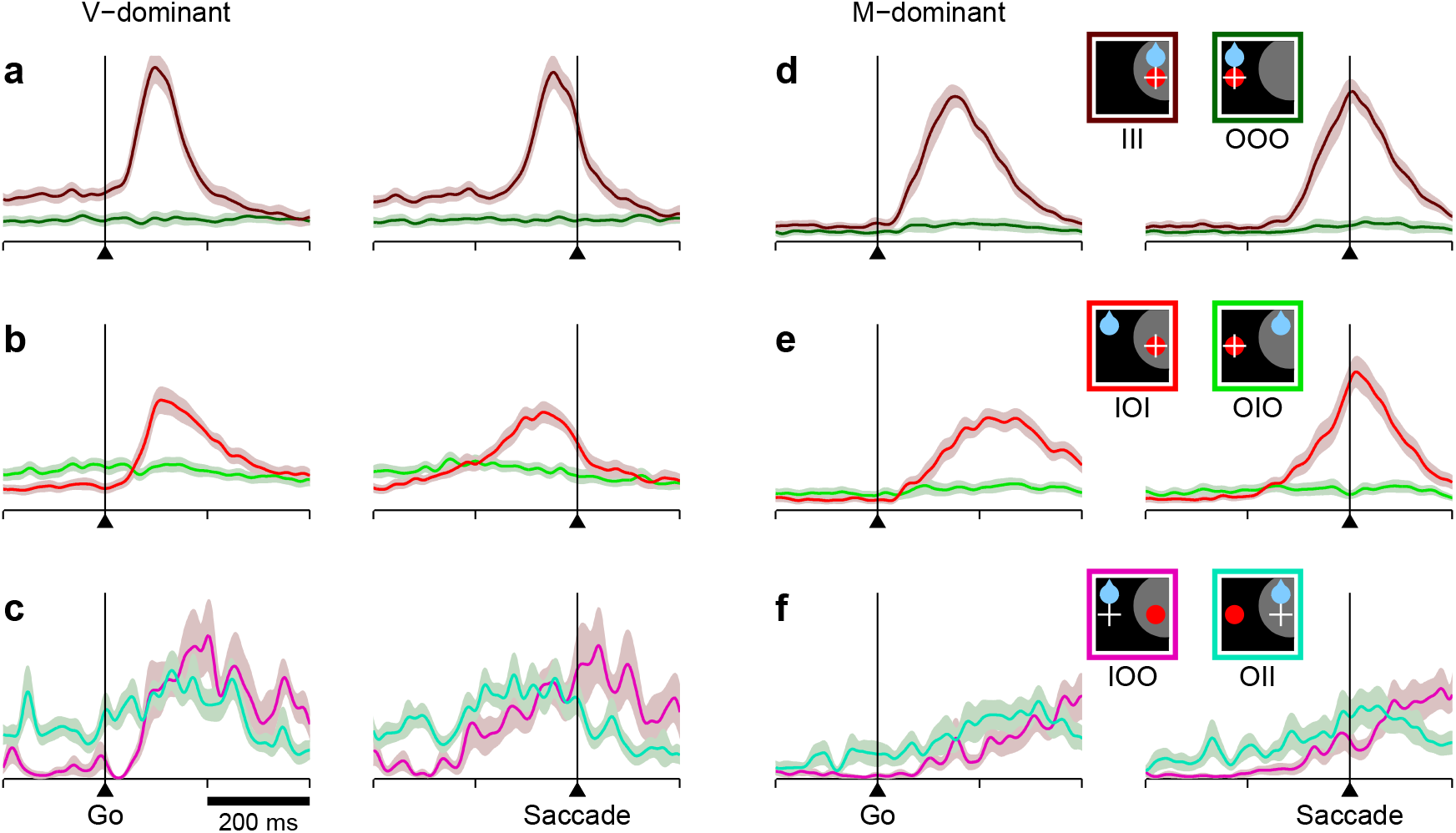
Responses of predominantly visual and predominantly movement-related neurons in FEF across conditions and outcomes. Each panel shows normalized firing rate as a function of time for different combinations of target (red circle), reward (blue drop), and saccade (white cross) locations relative to the RF (gray area), as indicated by the icons on the right. **a-c**, Results are as in Fig. 3a-c, except based on the 33% of the cells (*n* = 21) that were most visual, i.e., had the lowest visuomotor index values. **d-f**, As in panels **a-c**, except based on the 33% of the cells (*n* = 21) that were most motor, i.e., had the highest visuomotor index values. Note differences in timing between visual and movement-related traces, but similar relationships between motor plans into and away from the RF across conditions.

**Supplementary Figure 3.**
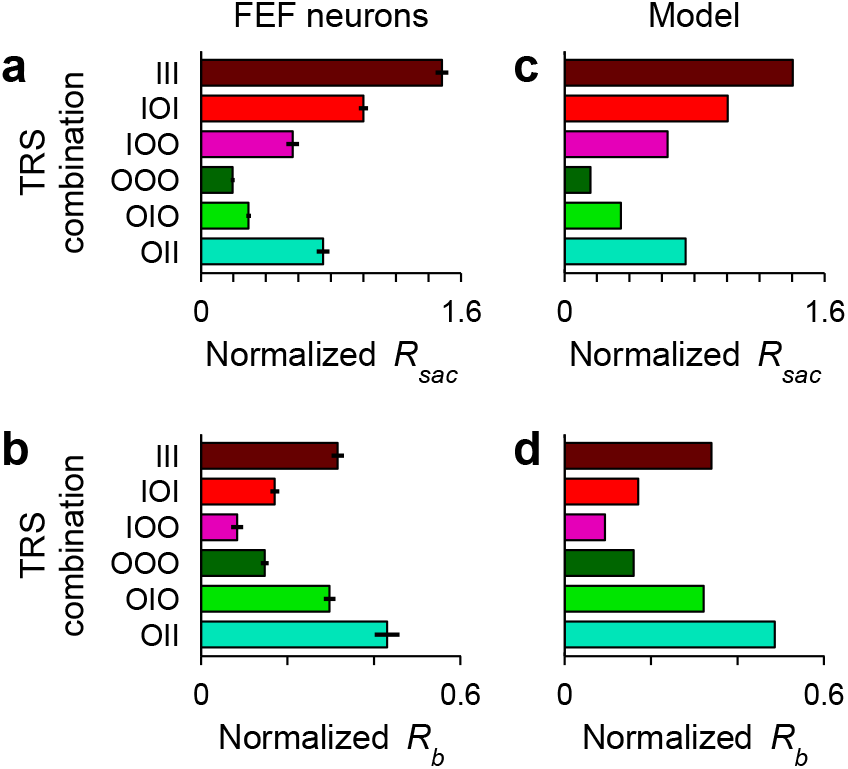
Variations in baseline and threshold levels in FEF and in the saccadic competition model. **a**, Mean normalized activity preceding saccade onset, *R_sac_*, for each target/reward/saccade (TRS) combination. Data are from the same population and in the same format as in Fig. 3d, but here activity is based on the firing rate of each neuron computed in a fixed 50 ms window ending at saccade onset. Also, each unit's responses were normalized before averaging across cells. Error bars indicate ±1 SE across cells. **b**, Mean baseline activity (preceding the go signal), *R_b_*, for each experimental condition. Data are from the same population and in the same format as in Fig. 3e, but here each unit's responses were normalized by the same factor as in **a** before averaging across cells. **c, d**, As in **a** and **b**, but from model simulations; same runs as in Fig. 4d-f. Presaccadic activity, *R_sac_*, was equated to the response at threshold crossing. Baseline activity, *R_b_*, was equated to the response at the go signal. To compare the FEF and model responses in the same scale, both data sets were rescaled so that *R_sac_* in the IOI condition was equal to 1.

**Supplementary Figure 4.**
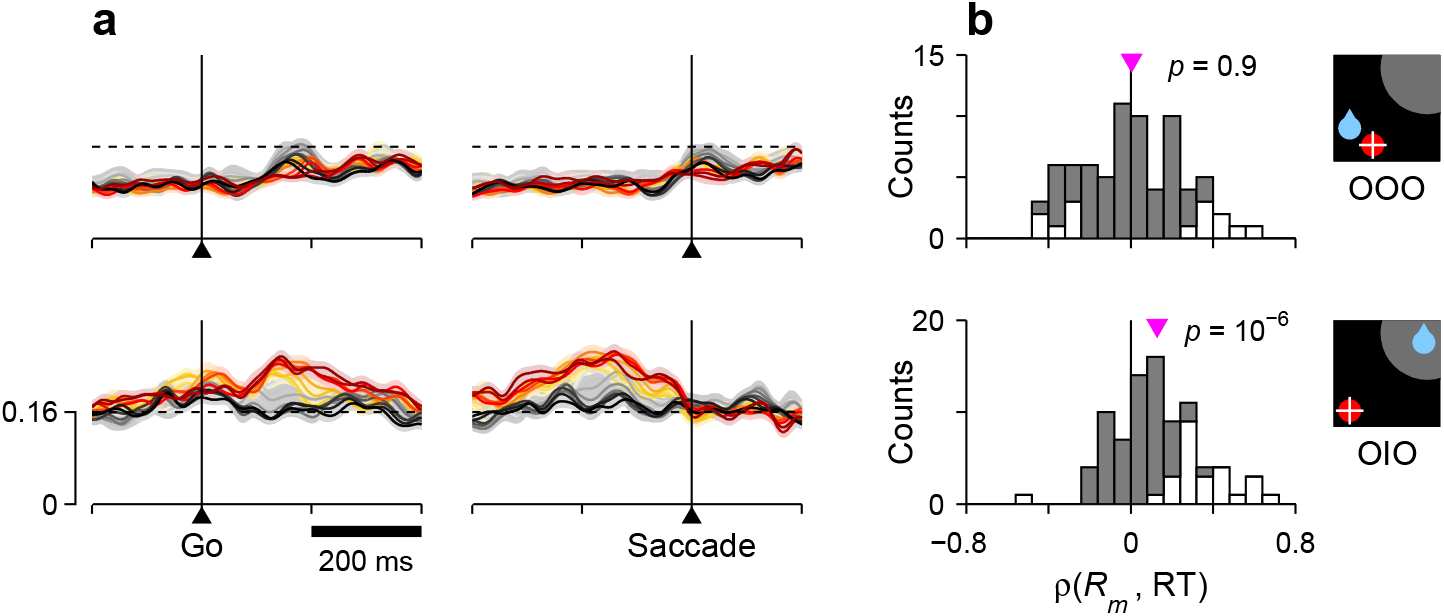
Correlation between RT and activity during correct saccades away from the RF. For each row, the corresponding target/reward/saccade configuration is depicted by the icon on the far right. **a**, Normalized neural responses. Each colored trace corresponds to population firing rate as a function of time for a subset of trials around a particular RT quantile. Traces are the same as in Fig. 6f, h, except with a shorter scale on the y axes, as marked on the bottom panel. The dashed reference line is identical across panels. **b**, Distributions of Spearman correlation coefficients between RT and mean activity, *ρ*(*R_m_*, RT), where *R_m_* is the mean response between the go signal and saccade onset. The data in each histogram are from the same 84 cells used in Fig. 6. White bars correspond to significant correlations (*p* < 0.05). Pink triangles mark mean values, with significance from signed-rank tests shown next to them.

